# Longitudinal Microbiome-based Interpretable Machine Learning for Identification of Time-Varying Biomarkers in Early Prediction of Disease Outcomes

**DOI:** 10.1101/2024.10.18.619118

**Authors:** Yifan Dai, Yunzhi Qian, Yixiang Qu, Wyliena Guan, Jialiu Xie, Duan Wang, Catherine Butler, Stuart Dashper, Ian Carroll, Kimon Divaris, Yufeng Liu, Di Wu

**Affiliations:** Department of Biostatistics, Gillings School of Global Public Health at University of North Carolina at Chapel Hill; Department of Nutrition, Gillings School of Global Public Health at University of North Carolina at Chapel Hill; North Carolina School of Science and Mathematics; Melbourne Dental School, University of Melbourne; Department of Pediatric Dentistry and Dental Public Health, Adams School of Dentistry, University of North Carolina at Chapel Hill; Department of Epidemiology, Gillings School of Global Public Health, University of North Carolina at Chapel Hill; Department of Statistics and Operations Research, University of North Carolina at Chapel Hill; Department of Genetics, School of Medicine, University of North Carolina at Chapel Hill; Department of Biomedical Sciences, Adams School of Dentistry, University of North Carolina at Chapel Hill

## Abstract

Information generated from longitudinally-sampled microbial data has the potential to illuminate important aspects of development and progression for many human conditions and diseases. Identifying microbial biomarkers and their time-varying effects can not only advance our understanding of pathogenetic mechanisms, but also facilitate early diagnosis and guide optimal timing of interventions. However, longitudinal predictive modeling of highly noisy and dynamic microbial data (e.g., metagenomics) poses analytical challenges. To overcome these challenges, we introduce a robust and interpretable machine-learning-based longitudinal microbiome analysis framework, LP-Micro, that encompasses: (i) longitudinal microbial feature screening via a polynomial group lasso, (ii) disease outcome prediction implemented via machine learning methods (e.g., XGBoost, deep neural networks), and (iii) interpretable association testing between time points, microbial features, and disease outcomes via permutation feature importance. We demonstrate in simulations that LP-Micro can not only identify incident disease-related microbiome taxa but also offers improved prediction accuracy compared to existing approaches. Applications of LP-Micro in two longitudinal microbiome studies with clinical outcomes of childhood dental disease and weight loss following bariatric surgery yield consistently high prediction accuracy. The identified critical early predictive time points are informative and aligned with clinical expectations.

## Introduction

Microbial communities within the human body are highly dynamic, continuously adapting in response to a complex interplay of host and environmental factors [1– 4]. These communities are essential for maintaining host health, modulating immune function, and facilitating metabolic processes [5, 6]. Studies have revealed that the composition and function of the microbiome can shift significantly over time, impacted by dietary changes, antibiotic exposure, lifestyle alterations, and disease states [7– 9]. The temporal fluctuations in microbial communities highlight the microbiome’s responsiveness and potential role as an indicator of health and disease [10].

The temporal development of the microbiome has therefore become an important topic of microbiome research, and longitudinal human microbiome studies are becoming increasingly available [4, 11, 12]. Capturing the dynamics of microbial communities over time is not only informative from a microbiological standpoint – it can provide insights into microbial events preceding disease development, progression, and treatment outcomes. Such information is relevant to many human conditions and diseases including cancers and inflammatory diseases [13–16]. The temporal development of the microbiome in various niches across the lifecourse may reflect endpoints of complex interactions between human behaviors (e.g., nutrition), innate biology, and the environment [11, 17, 18]. Understanding longitudinal interactions between the microbiome and the host is crucial for developing precise diagnoses, prognoses, and underlying mechanisms. Consequently, predicting incident disease outcomes using microbiome data is a compelling reason to invest efforts in the study of time-sensitive disease biomarkers with the potential for clinical translation.

Leveraging longitudinal microbiome data to predict disease outcomes is an ambitious goal, as it demands models that can account for the high-dimensional, sparse, and often non-linear nature of microbiome datasets. Traditional machine learning (ML) algorithms, such as support vector machine (SVM, [19, 20]), random forest (RF, [21]), and XGBoost [22], have shown promise in cross-sectional studies by identifying microbial markers of health and disease states. [23–25]. Meanwhile, deep learning (DL) methods, including recurrent neural networks (RNN, [26]), long short-term memory networks (LSTM, [27]), and gated recurrent units (GRU, [28]), offer potential by modeling these temporal relationships and capturing the changing microbial landscape with each time point.

Despite recent advances in predictive modeling, a key challenge in leveraging microbiome data for disease prediction lies in selecting the most informative microbial taxa as inputs, especially in longitudinal studies where data complexity grows. Longitudinal microbiome datasets are notoriously sparse and high-dimensional, often containing hundreds or thousands of microbial taxa, yet they are typically constrained by small sample sizes [29]. Standard feature screening, such as least absolute shrinkage and selection operator (lasso, [30, 31]) and sparse partial least squares (sPLS, [32]), generally assume linear microbial effects and may therefore miss non-linear microbial signals. Moreover, these traditional methods struggle in longitudinal setups, where preserving the continuity of microbial changes across time points is essential for capturing the trajectory of microbial effects. Methods like lasso and sPLS may select microbial features at isolated time points, often breaking the longitudinal data structure and hindering analysis of microbial dynamics over time. Advanced feature selection for non-linear microbial effects and longitudinal microbiome data is needed to achieve improved prediction power and capture the time-varying pattern.

Another key research question lies in identifying the time intervals during which signals from the microbial community are strongest for predicting disease onset or progression, especially when detailed mechanistic information is unavailable. This question can be answered by examining individual time points or by aggregating insights over multiple intervals. Suppose that the longitudinal microbial data are collected at times *t*_1_, …, *t*_*i*_, …, *t*_*q*_, and the outcome (i.e., disease endpoint) is observed at a later time point *t*_*r*_. One approach is to use the longitudinal data as a reference to identify the most important single time point for prediction, as the *visit-wise* prediction, by using microbiome data at each time point *t*_*i*_ as predictors independent of other time points. The other approach is to identify the earliest time points where predictive information is sufficient, beyond which additional time points may not further improve performance, as the *cumulative* prediction, by using the microbiome data corresponding to time points from *t*_1_ to *t*_*i*_, up to *t*_*q*_. While visit-wise approaches provide snapshots of microbial states at discrete intervals, cumulative prediction enables a more holistic understanding of the microbial trajectory leading up to a health event. In most applied settings—including translational research, biomarker discovery, or clinical practice—the interpretability of predictive models is crucial. The key objective is to understand how individual microbiome taxa interacts with the disease progression over time. To enable interpretable feature identification, common ML/DL-based methods, such as SHAP values [33–35], provide insights into feature importance but are less computationally efficient and lack formal significance testing. Alternatively, methods with formal statistical testing such as knockoffs [36, 37] and hold-out randomized tests (HRT, [38]) involve sampling from feature distributions with parametric covariate assumptions that are not suitable for highly skewed, zero-inflated, and high dimensional microbial data [29, 39]. Recent methods that sample from feature permutations [40, 41], such as PermFit [42] which is an ML/DL-based feature importance test for high-dimensional data, are superior in accommodating the skewed distribution of microbial data and avoid strong parametric assumptions. While enjoying the flexibility of these non-parametric tests, they cannot be directly applied to high-dimensional longitudinal data due to their reliance on the performance of ML models, which can be unstable for interpretation in this case.

Here, we introduce LP-Micro, i.e., Longitudinal Prediction Microbiome model, a comprehensive framework that integrates microbial feature screening, predictive modeling, and biomarker identification specifically tailored for longitudinal microbiome data. LP-Micro incorporates DL methods with ensemble learning [43, 44] alongside traditional ML models to build a predictive model well-suited for longitudinal microbiome data. In that, we propose the use of polynomial group lasso for feature selection in LP-Micro. LP-Micro first uses a polynomial spline to approximate the non-linear additive effects [45–47] and then groups the polynomial effects of a microbial feature (e.g., one taxon) across all time points (e.g., study visits) as one group [48–51], ensuring simultaneous selection of all features in the same group. Effectively, polynomial group lasso enables the screening of complete trajectories of informative microbial features (e.g., taxa) for downstream analyses. To allow for interpretable pre-screened features previously selected by group lasso, we propose to generalize PermFit [42] for longitudinal data where group testing is needed, by extending the original inference from testing a single feature to testing a group of features. This step also provides *p*-values for further interpretation, which the standard ML/DL methods can’t provide. LP-Micro results in superior prediction accuracy in both simulation and in two longitudinal clinical microbiome datasets, one for early childhood caries (ECC) and the second for weight loss following Bariatric Surgery (BS). In this paper, we demonstrate that the LP-Micro framework is useful for the identification of critical time points and microbial features. LP-Micro can be generalized for use with longitudinal high-dimensional data to jointly predict later outcomes and generate interpretable predictive features.

## Results

### Overview of LP-Micro

An overview of the analysis framework in LP-Micro is presented in Figure 1. Briefly, in the context of longitudinal studies, LP-Micro leverages microbial data from early time points to predict later outcomes, i.e., incident health and disease statuses. As mentioned above, with LP-Micro we are poised to achieve cumulative prediction using longitudinal microbiome data.

**Fig. 1:**
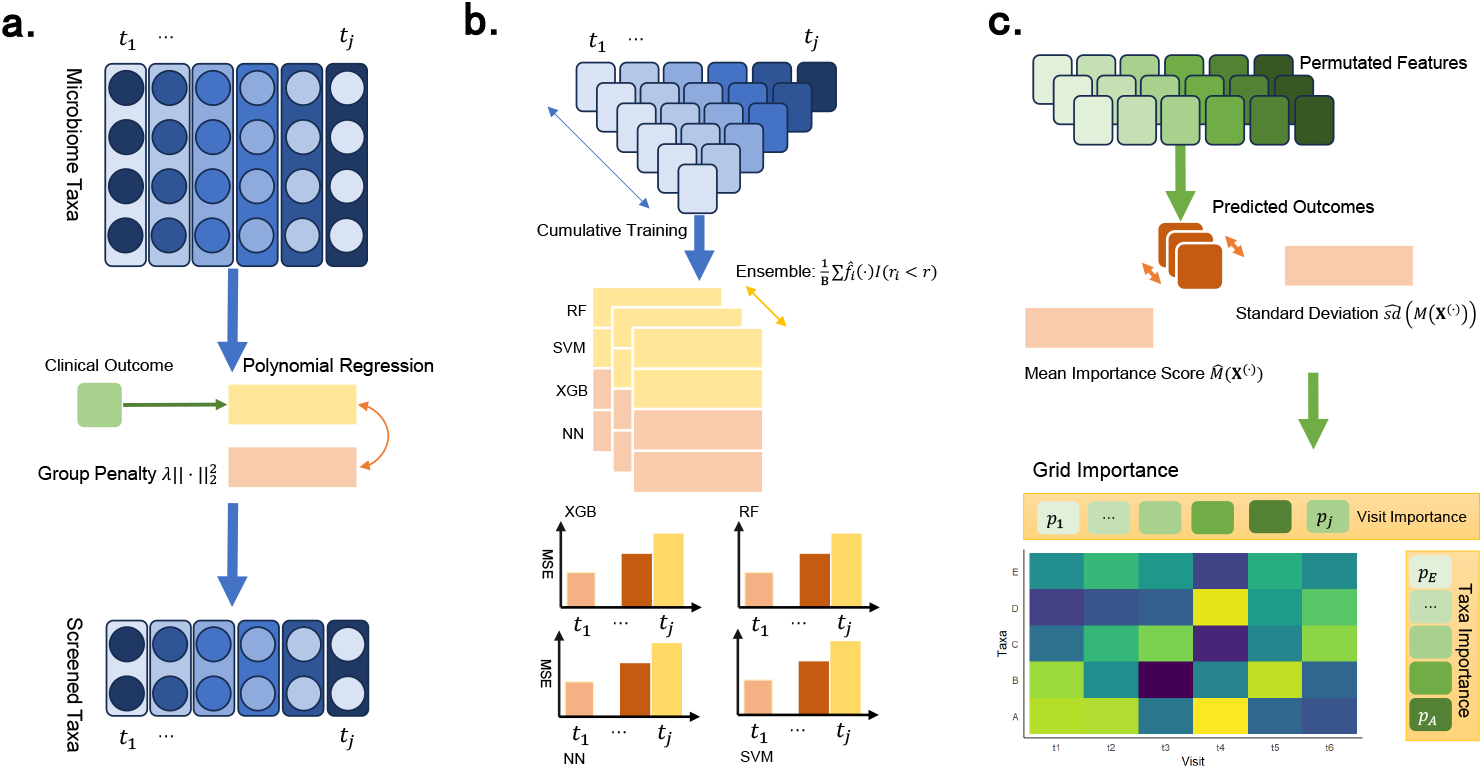
Flow chart of LP-Micro. The LP-Micro framework consists of three steps: **(a)** Pre-screening of microbial taxa using polynomial group lasso. **(b)** Cumulatively training and ensembling ML algorithms. **(c)** Prediction of the clinical outcome and interpretation of the contribution of microbiome taxa and time points.

There are three embedded steps in the development of LP-Micro. Firstly, to reduce the high data dimensionality, LP-Micro screens important microbial taxa with polynomial group lasso. To select a candidate set of most disease-predictive taxa, we extend group lasso to polynomial group lasso, allowing for nonlinear relationships between taxa and outcomes. According to previous work [46], feature groups can be defined by including one taxon derived from the feature-wise natural cubic spline method at each time point. Each group of features represents a taxon at one time point, which will be selected or discarded by its predictive ability. For cumulative prediction, we gather the above polynomial variables at each time point so that this one group of features represents the taxon in both nonlinear and longitudinal terms. In other words, each group of features represents a taxon across multiple time points. The selected group indicates that the abundance trajectory of the taxon is predictive of later disease outcomes.

Secondly, LP-Micro leverages the screened subset of microbial taxa to predict the clinical endpoints of interest. In this step, we deploy a variety of ML models: lasso, XGBoost, RF, SVM, as well as four deep learning architectures: fully-connected neural networks (NN), LSTM, GRU, and CNN-GRU. Importantly, we employ ensemble learning [43] to stabilize the performance of these deep learning methods. The effectiveness of these models is evaluated on a testing dataset, with raw models (i.e., those without feature screening and ensemble enhancements) serving as benchmarks for comparison. Using LP-micro, the optimal ML prediction model and the most relevant time points or periods are identified and reported.

Thirdly for interpretation, upon identifying the best prediction models, LP-Micro finally calculates permutation importance scores and corresponding *p*-values to quantify individual feature (i.e., microbial taxon) effects. Additionally, it assesses feature importance by grouping data in two ways: (i) by aggregating all time points for each taxon to determine taxon-wise importance across the study, and (ii) by aggregating all selected taxa at a single time point to evaluate the importance of specific time points (e.g., study visits) for so-called visit-wise importance score. To clarify, the visit-wise importance score here in LP-Micro is based on a likelihood function allowing all time points of data for cumulative prediction (details in Methods), and it does not necessarily correspond to visit-wise prediction, wherein data from only one time point are used. The visit-wise importance score in LP-Micro suggests which time points shall be least missed to achieve the best cumulative prediction results, instead of answering which single time point is most important. Therefore, this approach enables the reporting of both the most disease-relevant microbial taxa and the most critical time points in a period of time for predicting host development or disease progression.

In summary, compared to existing ML methods for disease prediction, LP-Micro offers three key advantages: (i) effective utilization of longitudinal microbial data; (ii) rigorous feature screening; (iii) generation of *p*-values for both temporal and single time point microbial effects, as well as the importance of time. In this paper, we evaluate the performance of LP-Micro through comprehensive simulation studies and application to two longitudinal clinical datasets: childhood dental disease (early childhood caries, ECC) data generated in the VicGen cohort study [52, 53] and weight loss data after bariatric surgery (BS) [54].

### LP-Micro performance evaluation in simulations

We evaluated the performance of LP-Micro in specifically simulated longitudinal microbiome data. Because sample sizes of this type of data are typically modest compared to other scenarios where machine learning is commonly used, we simulated microbiome data including 100-500 taxa from 120 participants measured at five time points. All simulated study participants have data collected at the same five time points. We generated five causal features (i.e., microbial taxa) that are associated with the outcome (i.e., disease), among totals of 100, 200, or 500 simulated microbial taxa, thus introducing three levels of signal sparsity (i.e., 5%, 2.5%, 1%). A continuous disease outcome was simulated. To incorporate time-varying microbial effects, one of the five causal taxa is associated with disease across all five visits, while the other four are only associated with disease in the last two visits. Consequently, the outcome variable is simulated based on a model including these 13 total microbial covariates (i.e., features in the prediction model). The identification of the correct causal taxa/covariates among a varying number of total simulated taxa is therefore the evaluation of feature-wise importance detection. The details of the complex, time-varying effects inspired by real data are described in the Methods section.

In this simulation, we attempt to predict the disease outcome using all time points. Although our focus is on cumulative prediction, evaluating predictions across all time points in this simulation can reflect cumulative prediction evaluations at multiple time points in real-world scenarios. To evaluate the effectiveness of LP-Micro under these simulated scenarios, we assess the pipeline’s performance based on three criteria that correspond to the three steps in LP-Micro: (i) pre-screening efficacy: the number of causal covariates selected by the pre-screening using polynomial group lasso, (ii) prediction accuracy: the prediction error of ML models, (iii) feature importance: the effect importance score assigned to the causal microbial features.

We begin by comparing the pre-screening efficacy of polynomial group lasso with standard lasso and sPLS [32, 55] considering the number of selected causal covariates. The results indicate that LP-Micro provides a refined feature set of microbial taxa for downstream analysis (Table 1). Specifically, the pre-screening step of LP-Micro using polynomial group lasso selects fewer than 50 variables and maintains on average more than 10 important variables (i.e., out of 13 with causal effects). Importantly, the number of correctly selected causal variables remains stable while the data dimensions increase (Table 1). In contrast, when faced with 500 simulated microbial taxa, the average number of causal covariates selected by the standard lasso is as low as 5.3. Similarly, sPLS, previously used to select important microbial taxa contributing to the temporal development of ECC [12], fails to select the majority of causal microbial variables when the effect is nonlinear and complex. The average number of selected causal variables is 5.50 in the context of 500 simulated microbial taxa.

**Table 1:**
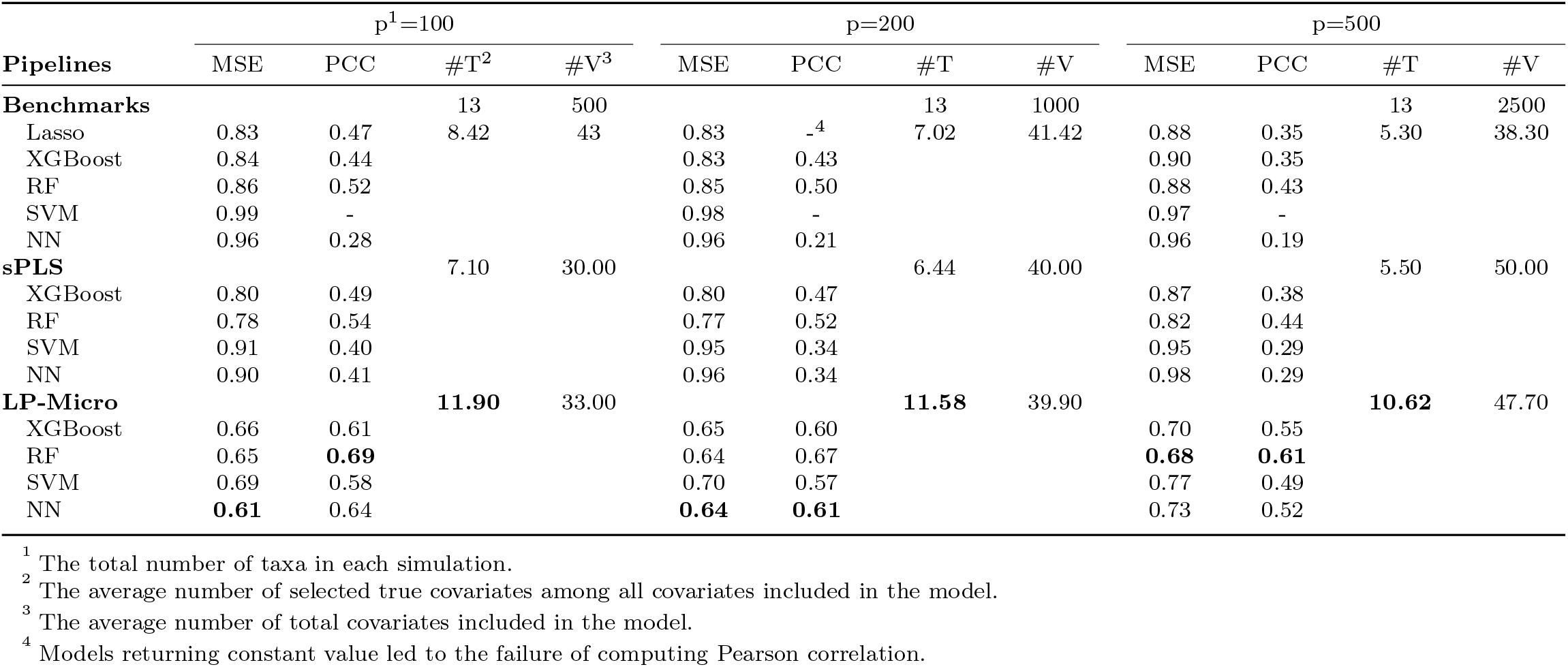
Prediction results in simulations. We compared the performance of machine learning models using (a) all features, denoted as benchmarks, (b) features selected by sPLS, (c) features selected by LP-Micro.

Next, we compare the prediction performance of LP-Micro ML models to (i) corresponding benchmark models of XGBoost, RF, SVM, and NN with complete features as input, (ii) models with screened input features by sPLS, using mean squared error (MSE) and Pearson correlation (PCC) between predicted and observed outcomes as metrics. As shown in Table 1, all other benchmarked ML models have larger prediction errors than lasso regression. However, with LP-Micro screening and ensembling, using a simple NN results in prediction MSE as low as 0.61, outperforming other methods. XGBoost, RF, and SVM also achieve more accurate prediction performances compared to their corresponding benchmarked models in this simulation setting. In comparison, while sPLS also improves the performance of ML models by reducing covariate complexity, the extent of improvement from sPLS is inferior to that of LP-Micro. For example, when faced with the scenario of 500 simulated microbial taxa, sPLS improves the prediction PCC of RF by only 1%, whereas LP-Micro increases it by 18%. We carried out similar comparisons for other NN architectures, including LSTM, GRU, and CNN-GRU (Supplementary Table 1) demonstrating that none of the advanced architectures result in significant improvement over the simple NN. Some architectures, such as LSTM, even experience overfitting, wherein the predictive error on the validation data increases as training progresses.

Lastly, we compare feature importance scores of LP-Micro models to those obtained from benchmark models using PermFit. Recall that the data-generating process assumed that one microbial taxon has a consistent effect over all five time points, four microbial taxa have effects only in the last two time points, and the remaining taxa are not associated with the disease outcome. LP-Micro accurately identified the patterns of longitudinal microbial effects on the outcome, highlighting the importance of the fourth and fifth visits (Figure 2a). Moreover, it detected the one microbial taxon with the simulated constant temporal effect starting from the first visit. In contrast, PermFit based on RF, SVM and NNs missed the temporal microbial effects. Although PermFit based on XGBoost correctly captured the pattern of time-varying microbial effects, its importance scores were lower than those from LP-Micro, indicating the efficiency benefit brought about by feature screening. In terms of microbial biomarker identification, the importance scores from LP-Micro align well with the simulation settings, assigning high scores to the causal microbial taxa (Figure 2b). Conversely, PermFit failed to identify those causal microbial taxa and instead highlighted non-causal features as important. For further comparison, we implemented HRT (which samples the microbial distributions using Gaussian distributions instead of permutations) as a means of deriving feature importance scores from LP-Micro and the benchmark models. The inferior performance of HRT compared to both PermFit and LP-Micro, underscored that the Gaussian sampling used by HRT is unsuitable for microbial data as HRT has much worse performance than LP-Micro to identify the true signals of important predictive features (Supplementary Figure 1).

**Fig. 2:**
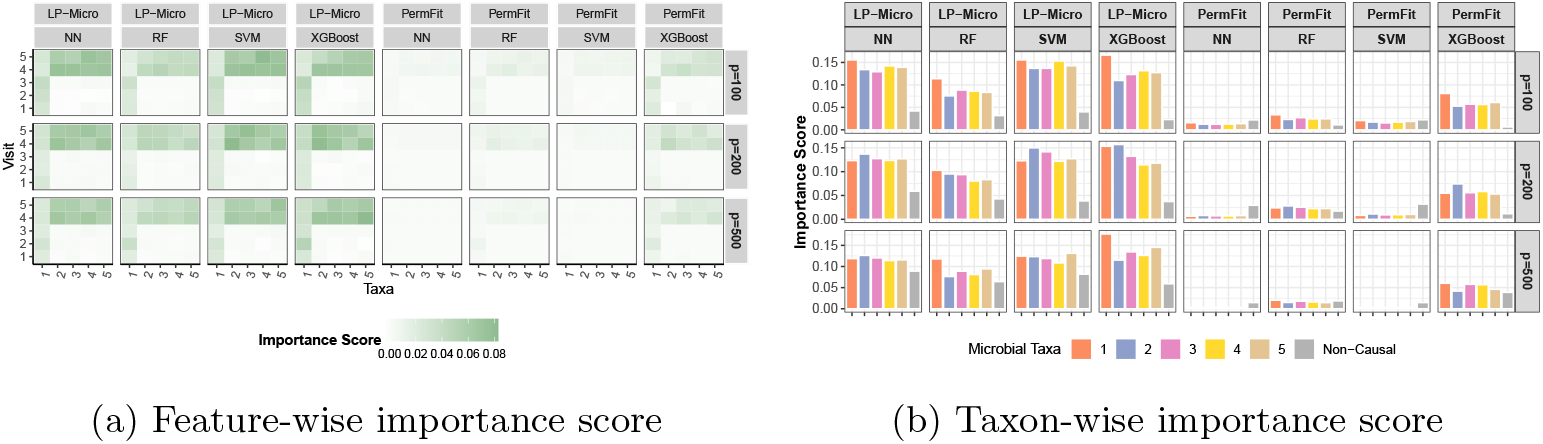
Feature importance in simulations. **(a) Microbial feature importance at each time point.** The x-axis represents five causal microbial taxa, and the y-axis represents each of five visits. Each grid represents the feature importance of the microbial taxa during the corresponding visit row. **(b) Overall microbial feature importance across time points**. The x-axis consists of five causal microbial taxa and the set of non-causal microbial taxa. The y-axis represents the taxon-wise importance across five visits, obtained by aggregating importance scores in (a). Here, *p* is the total number of taxa in each simulation.

### Application of LP-Micro to study early childhood caries (ECC) in the VicGen cohort study

The VicGen cohort consists of compositional microbial profiles generated for 134 children at six time points, with a binary disease outcome indicating the presence of dental caries in early childhood (ECC, i.e., cavities in primary teeth) measured via a clinical examination at age five [12], defined as having an International Caries Detection and Assessment System (ICDAS) score 3 or above [56]. This is the threshold of macroscopic enamel loss, i.e., a frank cavity which is an unambiguous and irreversible stage of dental caries, likely to be clinically consequential and commonly used in observational and experimental studies of childhood dental disease [57, 58]. The six time points of microbiome collection range from one or two months of age up to five years old and are generally matched across this cohort in this study (Figure 3a). We aim to predict ECC at age 5 using the longitudinal microbiome data of the first six time points up to 48 months of age. To address the importance of longitudinal microbial profiles in prediction, we reshape this aim as a cumulative prediction to understand up to which time points before 48 months old, the highest prediction accuracy and area under the curve (AUC) can be achieved. Because the ECC outcome at year five is known, multiple ML models in the prediction step of LP-Micro, i.e., RF, XGBoost, NN, and SVM, can be applied with multiple options at the feature selection step, i.e., baseline ML methods with either no specific feature selection, sPLS, PermFit, or group lasso in LP-Micro. We can also use the VicGen Study and the next weight change study to compare the performance of these models with varying feature selection options. We still include a simple model lasso for both feature selection and prediction as a more traditional reference model for comparison.

**Fig. 3:**
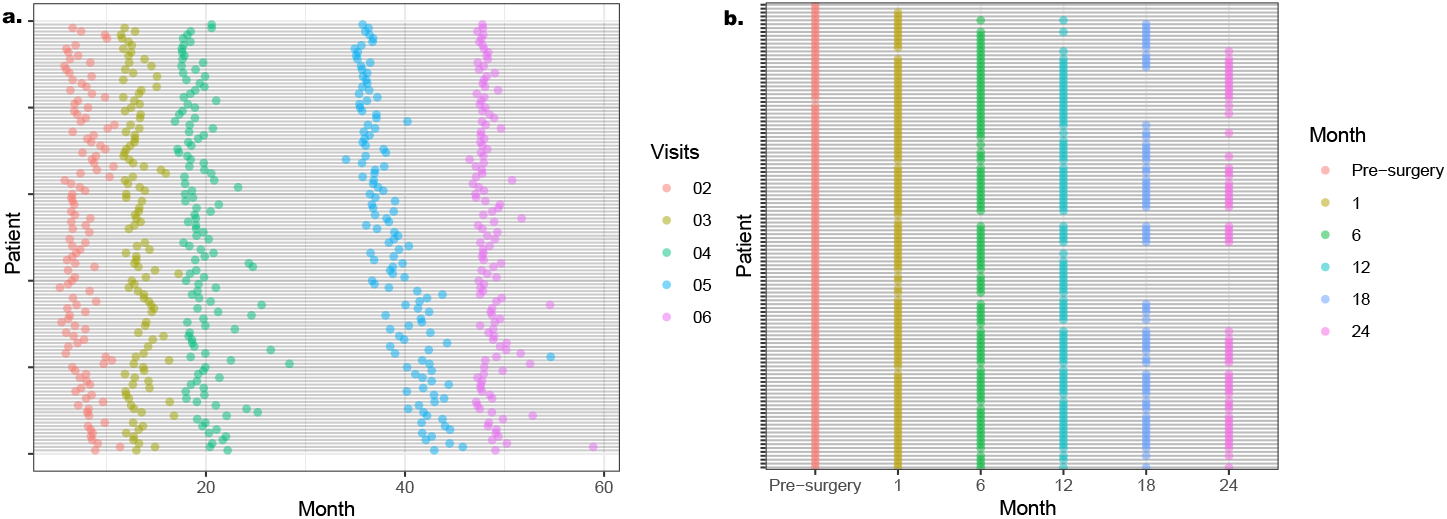
Timeline of longitudinal microbiome data collection in (a) the VicGen cohort investigating ECC, (b) the weight loss study following BS. The y-axis represents all available study participants from the two studies (*n* = 134 and *n* = 120, respectively), and the x-axis represents the time when their saliva or fecal samples were collected and analyzed. In the VicGen cohort, the exact timing of subjects’ first visits is not available, though the range of first visit times is detailed in the Methods section. Additionally, for 10 subjects in the VicGen cohort, the timeline is not shown due to missing recorded visit times.

The analysis for the VicGen cohort can be summarized as follows: First, we compared the prediction performance of the five models lasso, RF, XGBoost, NN, and SVM within LP-Micro, as well as to the five models across different feature selection options for evaluation of feature selection procedure (Figure 4a). Second, we used the two best performance ML models and the reference lasso model to conduct visit-wise prediction directly by using these existing models on each of the single time points, cumulative prediction by directly using these existing models on stacked data due to multiple time points, and the cumulative prediction using LP-Micro (Figure 5a). Third, we calculated the permutation feature importance scores for Operational Taxonomic Units (OTUs, equivalent to taxa, and mostly bacterial species in this study) and time points using the selected models (Figure 5b, 6). Essentially, this step quantifies the contribution of each microbial taxon at each time point to the cumulative prediction of ECC measured at age five, enhancing our understanding of their roles in ECC development during the early stage of life. Overall, the *p*-values of importance scores (Figure 6a) at variable-wise, visit-wise (Figure 5b) and taxon-wise (Figure 6b) were generated based on the full length of time points from first visit to the last visit at year five. We choose to use *p*-values of feature important scores instead of the scores in real data for interpretation because it is convenient to use *p*-values as a cutoff.

**Fig. 4:**
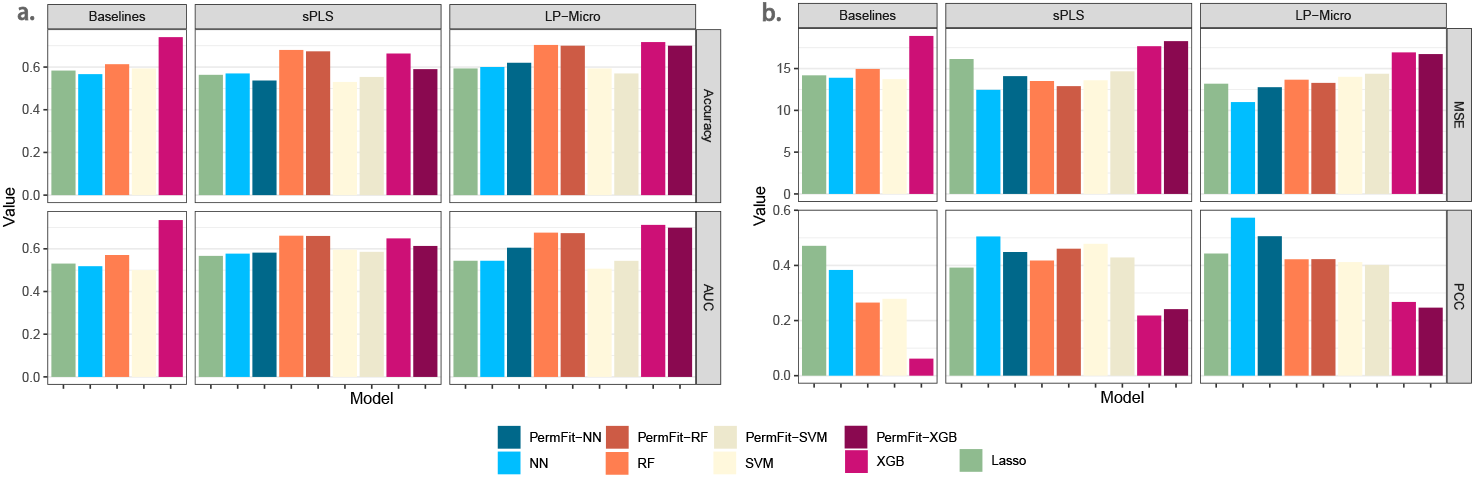
Prediction results to compare feature screen procedures for (a) the VicGen cohort investigating ECC, (b) the weight loss study following BS. Three sets of features are compared including first, the full longitudinal features combined by *p* taxa and *q* time points, second, the pre-screened features either by sPLS or by group lasso we developed in LP-Micro, and third, the selected features from the second sets further being narrowed down using the original PermFit. Selection of features in PermFit is based on the *p*-values of the permutation-based important scores per feature, so that the feature selection step and the interpretation step are the same step in PermFit. In the comparison, baseline models use full longitudinal features as input, while sPLS(-DA) and LP-Micro models use both the second and the third sets of features. In plots of LP-Micro, columns of PermFit-NN, PermFit-RF, PermFit-SVM, and PermFit-XGB further retrain LP-Micro models using only the corresponding second sets of features in LP-Micro with PermFit-generated p-values smaller than 0.1 so that the third set is a subset of the second set of features. Similarly, in plots of sPLS, columns of PermFit-Models also use only the corresponding second sets of features in sPLS with PermFit-generated p-values smaller than 0.1. In the VicGen study (a), only longitudinal microbial features were used for prediction, with no inclusion of demographic or clinical variables. Age was excluded as all participants were of similar age at enrollment. In contrast, for the weight loss study (b), both microbial features and five demographic variables were used for prediction, as detailed in Table 2.

**Fig. 5:**
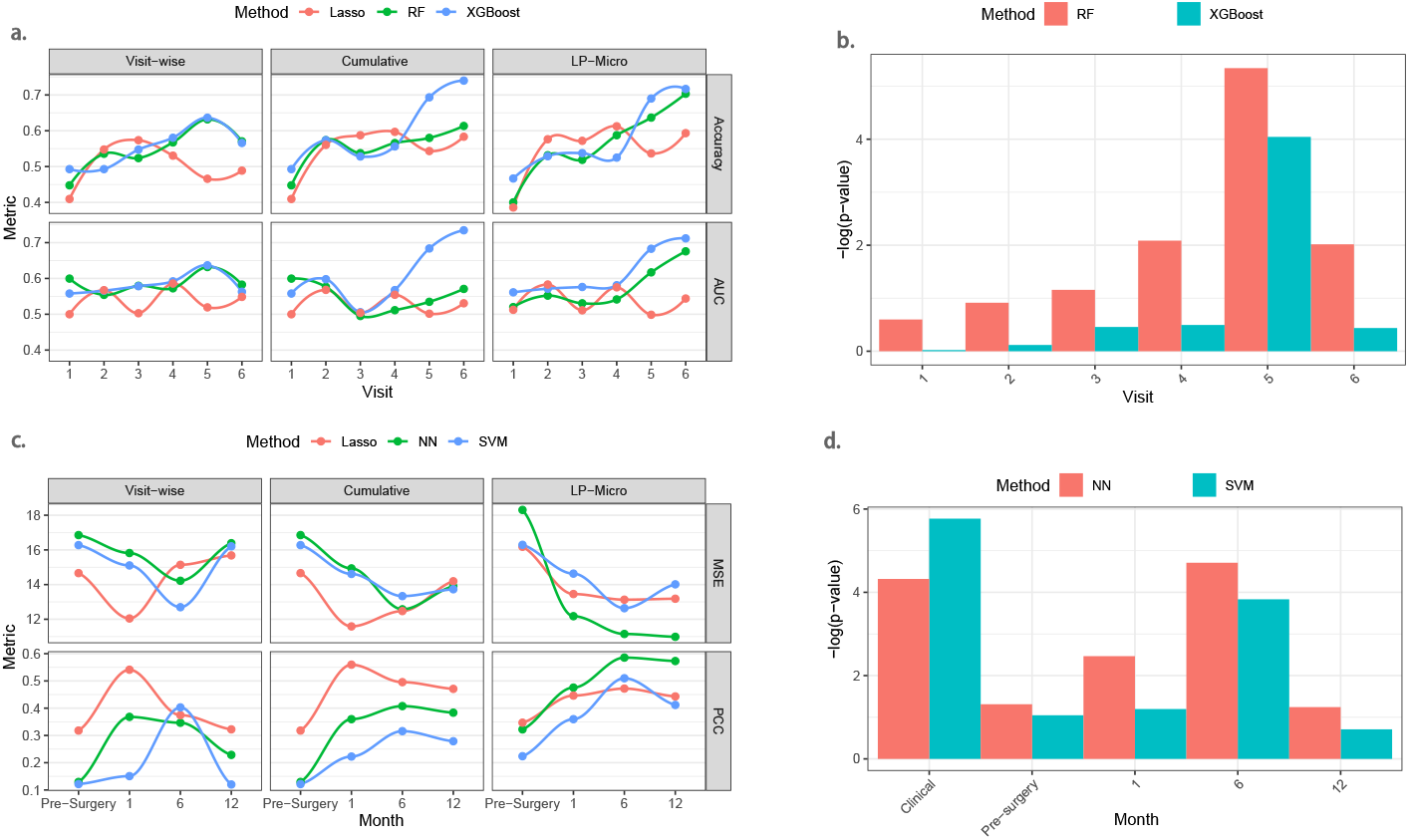
Results of visit-wise prediction, cumulative prediction, and LP-Micro. To accomplish visit-wise prediction and cumulative prediction, ML algorithms are trained using all microbial features. LP-Micro refines the cumulative prediction by microbial feature pre-screening and model ensembling. **(a) Prediction accuracy and AUC for ECC in the VicGen cohort**. The trajectory is smoothened using local polynomial regression. **(b) Visit-wise microbial importance scores calculated in LP-Micro for cumulative prediction of ECC using microbiome data up to the 6th visit. (c) Prediction MSE and PCC for BMI change in the weight loss cohort. (d) Visit-wise microbial** *p***-values of importance scores calculated in LP-Micro for cumulative prediction of BMI change prediction using microbiome data up to 12 months after BS**. In (b) and (d), log is 20 based. In the weight loss study (d), five covariates, i.e., height, sex, race, age, and surgery type (together as clinical in x-axis) with details in Table 2, were included in the prediction model. Pre-surgery in x-axis of (d) is the microbiome data before surgery.

**Fig. 6:**
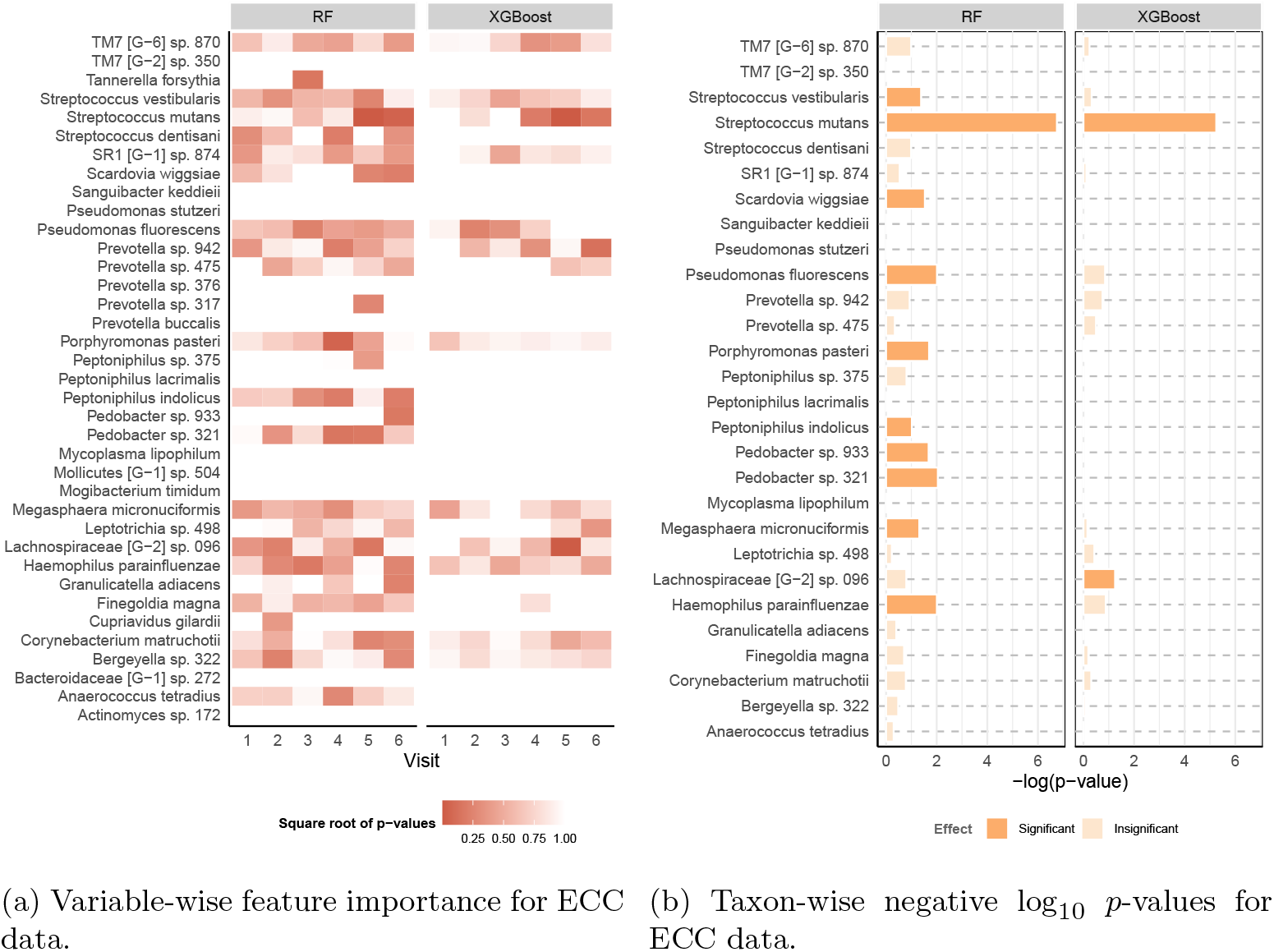
Feature importance for the ECC data. The y-axis consists of microbial taxa selected by LP-Micro. The *p*-value is computed on RF and XGBoost using microbial features up to the 6th visit. **(a) Microbial importance for ECC prediction at each visit**. Each grid represents the significance of the microbial taxon during the corresponding visit. Darker color indicates higher importance. **(b) Overall microbial importance across all visits for ECC prediction**. The x-axis represents the negative log_10_ *p*-values for each taxon in terms of their longitudinal effects on ECC. In (b), taxa are defined as significant if their permutation important scores have *p*-values smaller than 0.1.

### Feature Screening in LP-Micro improves ECC prediction

To validate taxa screening procedure by LP-Micro in prediction results, we compare the LP-Micro ML models with (i) the corresponding baseline models using full features as combination of taxa and time points, and (ii) the models using features screened by sPLS discriminative analysis (sPLS-DA). As shown in Figure 4a, LP-Micro improves the performance of most baseline models in terms of accuracy and AUC, with LP-Micro RF and LP-Micro XGBoost being top performers. XGB and RF have higher accuracy and larger AUC overall. Compared to other prediction frameworks, the performance of ML models with sPLS-DA screening is inferior to the corresponding algorithms of LP-Micro, indicating that LP-Micro selects more reliable microbial taxa for ECC prediction. Interestingly the baseline XGBoost also is among the best prediction model without our specific feature selection, however the feature importance scores in XGBoost, e.g., gain, do not have the corresponding *p*-values, the test for prediction is not at the level of grouped features for longitudinal data (e.g., multiple time points in one taxon) so that XGBoost provides less model interpretation in high-dimensional longitudinal prediction comparing to LP-Micro.

**Table 2:** Demographic and clinical characteristics of patients included in the weight change sub-cohort for our analysis (n=84).

| Variable | Number | Proportion (%) | Mean (± s.d.) |
| --- | --- | --- | --- |
| **BMI** | 84 |  |  |
| Pre-surgery | | | 45.64 (± 7.20) |
| 1 month | | | 40.27 (± 6.65) |
| 6 months | | | 34.28 (± 6.16) |
| 12 months | | | 32.71 (± 6.32) |
| **Age (years)** | 84 | | 42.63 (± 10.18) |
| **Height (inches)** | 84 | | 66.2 (± 3.36) |
| **Sex** |  |  |  |
| Male | 13 | 15.48 |  |
| Female | 71 | 82.80 |  |
| **Race/Ethnicity** |  |  |  |
| Caucasian | 60 | 71.43 |  |
| Black | 19 | 22.62 |  |
| Hispanic | 2 | 2.38 |  |
| Other | 3 | 3.57 |  |
| **Surgery** |  |  |  |
| RYGB | 54 | 64.29 |  |
| SG | 30 | 35.71 |  |

Furthermore, to validate the feature interpretation in LP-Micro, we assess the impact on model performance after selecting significant features with small *p*-values, following the original PermFit method [42]. A minimal performance change after feature selection indicates reliable feature interpretation, as most of the predictive information remains within the selected significant features. Specifically, we chose to retrain sPLS-DA models and LP-Micro models using significant features with *p*-values *<*0.1 in PermFit, denoting them as PermFit-ML models. Due to their high computational cost for high-dimensional data, PermFit models for the baseline algorithms failed to converge and are therefore unavailable. In LP-Micro and sPLS, the performance of PermFit-XGB and PermFit-RF, as shown in Figure 4a, remains either comparable or slightly lower to their corresponding models without PermFit. The stable performance upon feature selection suggests that the variance of ECC can mostly be explained by the microbial features identified by LP-Micro.

### Longitudinal microbial data boost ECC prediction

As the most important figure in this study, the prediction trajectory illustrates that the incorporation of longitudinal microbial features improves the prediction accuracy (Figure 5a). Although the improvement is limited at the initial visits, possibly due to weak microbial signals (i.e., the primary dentition may not be complete until the age of 2.5-3.0 years and the oral microbiome is still developing), incorporating previous microbial profiles improves the prediction accuracy starting at the 5th visit (39 ± 3.2 months-of-age). By the 6th visit (48.6 ± 1.6 months-of-age), the prediction accuracy of XGBoost incorporating information from previous visits is 74.0%, while the accuracy of visit-wise prediction is only 56.6% (Figure 5a). In contrast, RF, which tends to be more sensitive to high-dimensional data and prone to over-fitting, shows a more modest improvement. When past microbial profiles are concatenated directly, the accuracy increases from 57.0% to 61.3%. This smaller gain is likely due to the additional complexity introduced by the longitudinal data. Nevertheless, the feature selection process of LP-Micro mitigates this over-fitting issue, allowing RF to leverage the enriched information from longitudinal data more effectively. As a result, by the 6th visit, RF achieves a prediction accuracy of 70.0%.

LP-Micro provides insight to early changes in the oral microbiome that are predictive of later ECC development and may be reflective of early, sub-clinical changes in the oral ecology that could inform disease prevention, timely intervention, or disease management strategies. By the 5th visit at 39 months-of-age, cavities were clinically detectable in only 12 out of 65 children who were later diagnosed with ECC, with the remaining 53 children diagnosed at later stages, at 48 or 60 months of age. However, LP-Micro achieved a 70.0% accuracy in predicting the future development of cavities at this early stage (Figure 5a), suggesting that the salivary microbiome may be indicative of the disease process two years before frank cavities are macroscopically evident on children’s teeth.

The prediction curve of LP-Micro identified age three (i.e., the 39-month visit) as a critical time point for ECC prognosis using participating children’s salivary microbiome. As shown in Figure 5a, RF and XGBoost achieved the highest accuracy of visit-wise predictions at the 5th visit. The accuracy of longitudinal prediction substantially increases with the inclusion of microbiome information obtained at the 5th visit. Said differently, the 5th visit information boosts the prediction accuracy for ECC. The observation is consistent with previous findings reported from the same cohort, suggesting that the microbiome information from the initial visits is less associated with ECC than those of later visits [12]. In contrast, we noticed that baseline models like lasso do not produce desirable patterns. Group permutation tests further reinforce the observation by greater feature importance scores for microbiome profiles collected around 39 months-of-age compared to other visits. The microbial signal for ECC is the strongest at 39 months-of-age (Figure 5b). When the statistical significance level of the visit-wise importance scores (y axis in Figure 5b) calculated for cumulative prediction in LP-Micro using data from 1st time point to the 6th visit) is set to 0.05, RF model identifies that microbiome profiles from the 3rd to the 6th visits are significant, while XGBoost only identifies the 5th visit as the significant time point. Besides addressing the importance of microbial profiles from later stages, both RF and XGBoost also indicate that the microbial information before the 5th visit is increasingly important, whereas information from the 6th visit for ECC prediction is less important than the 5th visit. The trend in visit importance score implies that microbial effects on ECC are time-varying, potentially with a peak of effect size between the 5th and the 6th visits, providing additional insights compared to earlier reports [12].

### Interpretation of the identified microbial taxa is aligned with empirical evidence

The feature-wise permutation importance score identified significant microbial OTUs (i.e., bacterial taxa) associated with ECC development and highlighted *Streptococcus mutans* as the most predictive taxon (Figure 6). This is consistent with previous studies showing that *S. mutans* is highly correlated with dental caries [59–61]. Furthermore, we find that *S. mutans* begins to associate with ECC as early as at the 39 months-of-age visit, while the association is not significant earlier. Specifically, the *p*-values for *S. mutans* at the 5 and 6th visits from RF and XGBoost are smaller than 0.05. In contrast, the abundance level of *S. mutans* collected from other visits is less correlated with ECC. Our results also show that *S. mutans* is more predictive at 39 months-of-age (5th visit) than at the subsequent time point. The *p*-value of *S. mutans* at the 6th visit from RF is only slightly smaller than 0.05, while the *p*-value of the same OTU at the 5th visit is smaller than 10^−5^. This finding also helps explain the leap of longitudinal prediction trajectory at the 5th visit, implying that the new information brought by *S. mutans* at that time is crucial for timely disease diagnosis. This observation is also aligned with the expectation that microbiome changes should be detectable before the onset of clinically manifested disease.

Aside from *S. mutans*, RF and XGBoost also identify *Streptococcus vestibularis, Scardovia wiggsiae, Pseudomonas fluorescens, Porphyromonas pasteri, Peptoniphilus indolicus, Pedobacter* oral taxon 321, *Megasphaera micronuciformis, Lachnospiraceae*, and *Haemophilus parainfluenzae* as important predictors of ECC, and novel associations will need to be replicated and experimentally validated in future studies. Their longitudinal dynamics and temporal associations with ECC can be found in Supplementary Figure 2-4.

### Application of LP-Micro to study weight loss after Bariatric Surgery (BS)

This cohort study collected microbial data (1533 genera) and body mass index (BMI) measurements from 144 adult participants before their BS (pre-surgery as time 0) and 1, 6, 12, 18 and 24 months after surgery. Participants’ microbial profiles in the BS study were characterized through shotgun whole genome sequencing for time-series metagenomics data of 120 subjects (*n* = 120, [62]). Demographic variables, including participants’ height, sex, race, age, and surgery type are also available pre-surgery. Subjects have practically the same timeline of microbial sample collection (see Figure 3b).

BMI shows no significant change between 12 months to 24 months and more than 50% of participants were lost to follow up after month 12. Therefore, we only investigated prediction of BMI change between the 12th month post-surgery to that before surgery, using pre-existing demographic and clinical characteristics, such as the surgery type of BS, and longitudinal (0, 1, 6, 12 months) microbial profiles. Consequently, we included 84 individuals with longitudinal microbial data and no missing time points from pre-surgery to 12 months post-surgery, a total of four time points. The descriptive statistics of their demographic variables are summarized in Table 2.

### Feature Screening in LP-Micro improves predictability of weight loss

The genera selected by LP-Micro maintain most of the correlation with weight loss and lead to best prediction performance compared to baseline and models with sPLS screening (Figure 4b). LP-Micro NN outperforms all other predictive models. Moreover, LP-Micro enhances the performance of all tested ML models in terms of prediction MSE and PCC. In comparison, the performance of NN, XGBoost, and Lasso with sPLS is inferior to LP-Micro. These findings imply that LP-Micro selects reliable microbial taxa for weight prediction and feature interpretation. Furthermore, the performance of PermFit ML models, as shown in Figure 4b, is comparable to the baseline when using only significant features as predictors. The stable performance upon feature selection implies that obesity-related information is concentrated in the microbial genera selected by LP-Micro, resulting in reliable interpretation.

### Leveraging longitudinal microbial data improves prediction accuracy of weight change after BS

Incorporating microbial features obtained at multiple visits instead of only from one visit boosts the performance of ML models. This implies that earlier levels of microbial abundance are reflective and thus predictive of subsequent BMI changes. As shown in Figure 5c, the prediction of LP-Micro, which utilizes information from multiple visits, is more accurate than the visit-wise prediction in terms of MSE and PCC. However, we observe that the advantage of using longitudinal microbiome data diminishes when temporal microbial abundance is directly aggregated for training. For example, both NN and SVM show a marked decline in performance when longitudinal data is included. Their cumulative prediction accuracy does not significantly improve compared to visit-specific predictions. For instance, SVM achieves its highest accuracy (PCC *>* 0.4) using microbiome data from six months post-surgery (see the visit-specific plot in Figure 5c), whereas its cumulative prediction PCC remains below 0.35 (see the cumulative plot in Figure 5c). Additionally, the accuracy of both NN and SVM is consistently lower than that of lasso, for both visit-wise and cumulative predictions. In contrast, LP-Micro shows significant improvements in cumulative predictions that account for time-varying microbiome effects, delivering better results than visit-wise predictions. For example, at 6 and 12 months post-surgery, the prediction PCC of NN exceeds 0.55, outperforming other models. Interestingly, lasso performs best for early predictions of one-year BMI changes after BS, achieving the highest accuracy among models using only microbial abundance from one month post-surgery. However, its performance becomes inferior to NN and SVM as more time points are included. This aligns with our expectation, as LP-Micro has too few time points to be effective in one month after BS. Nevertheless, it still enhances the performance of NN and SVM in this case. Beyond the one-month mark, LP-Micro NN and SVM become stronger predictors of weight loss.

The pattern of prediction curves in Figure 5c suggests that microbial features after (versus before) surgery may be more important in predicting weight change. In the visit-wise prediction, lasso achieves its lowest prediction errors using information at one month after surgery, while NN and SVM achieve their best performance with microbial data collected at six months after surgery, suggesting that the gut microbial dynamics right after surgery may be most associated with the weight loss effect of BS. In contrast, the microbial profiles collected before surgery and 12 months after surgery, are shown to be less important in terms of accuracy, in all three prediction approaches. The group permutation test of LP-Micro further affirms our interpretation of time point importance (Figure 5d). Both SVM and NN identify clinical variables (including surgery type, race, gender, and age) and gut microbiome measured at 1 and 6 months after BS as important signs of future weight loss or weight gain, while later microbial profiles are less correlated with the weight change. Specifically, we find that pre-surgery clinical variables and microbial profiles from the first two visits post-surgery have significant effects on later weight gains (*p* ≤ 0.05), while microbiome information collected at the last visit is not important. Additionally, we re-conducted the analysis by removing the pre-surgery microbial features, and the results are similar to the analysis on the complete data as summarized in Supplementary Figures 5-7.

### LP-Micro provides insight in the temporal development of weight change after BS

According to permutation importance scores, age is one of the most important factors influencing the recurrence of obesity one year after BS (Figure 7). In contrast, gender and height appear not to be predictive of BS outcomes. This finding is similar to previous reports [63] showing that weight change after BS varies by age, regardless of gender and surgery type. Other demographic characteristics such as race are also significantly associated with patients’ BMI change. Furthermore, among the microbial genera selected by group lasso, *Acidilobus, Cloacibacterium, Cobetia, Escherichia, Litorilituus, Schaalia, Sulferhydrogenibium, Thermosulfuriphilus*, and *Turicibacter* showed significant effects in both NN and SVM in predicting patients’ weight gain (Figure 7b). Interestingly, as suggested by Figure 7a, we find that the effects of *Cobetia* and *Schaalia* are more significant before surgery or in one month after surgery, suggesting the potential of these two genera as early-stage biomarkers for weight loss. The longitudinal dynamics of those features and their time-varying association with post-surgery BMI change are summarized in Supplementary Figure 8-9. Notably, we find the correlation between most gut microbial taxa and the weight change non-linear.

**Fig. 7:**
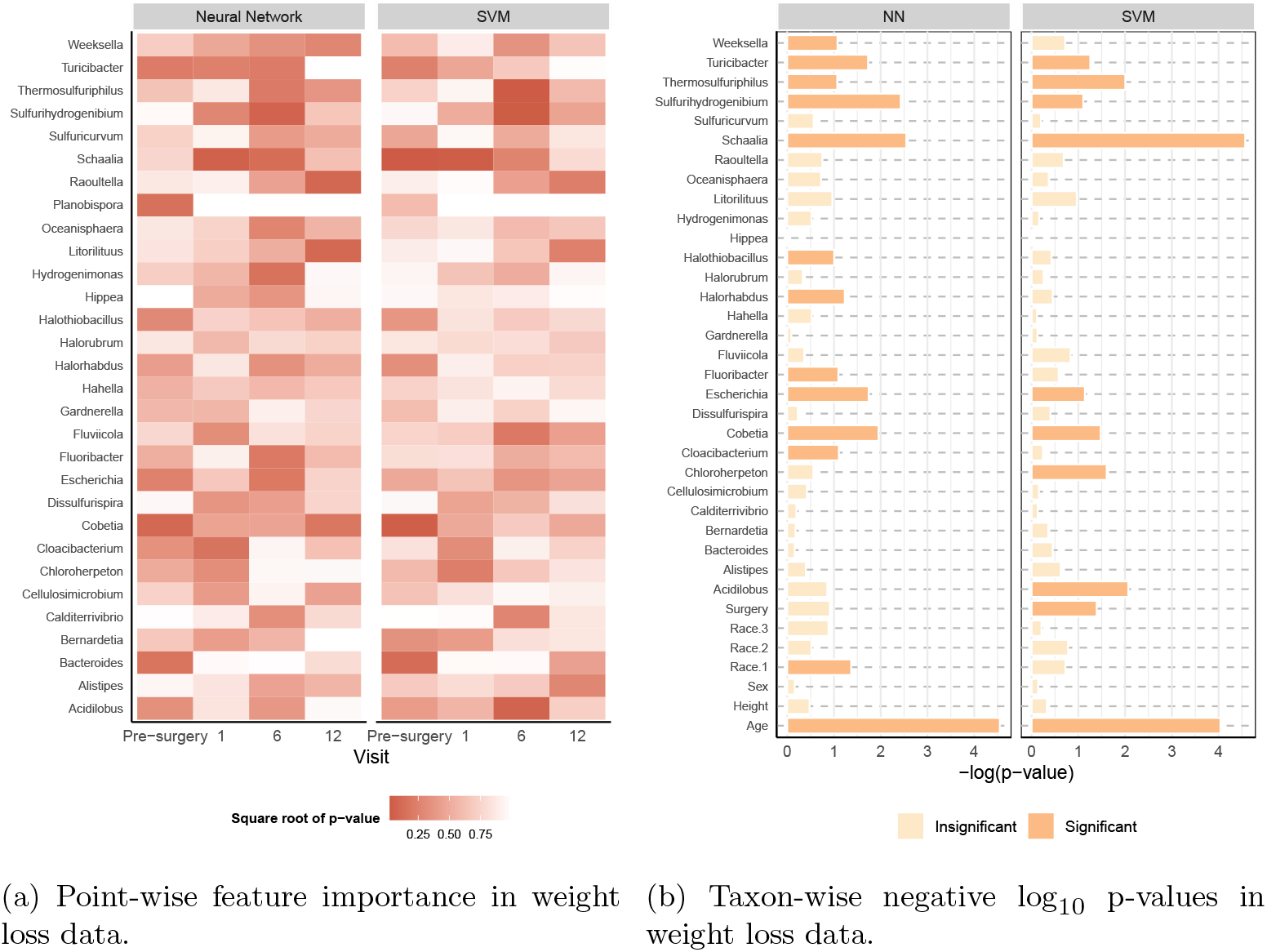
Feature importance for the weight loss data. The y-axis consists of microbiome genera selected by LP-Micro, and *p*-values are computed by PermFit. **(a)** Each grid represents the significance of the microbiome at each corresponding visit. Darker colors indicate higher importance. **(b)** The x-axis represents the log_10_ *p*-values for each taxon in terms of their longitudinal effects on BMI change.

## Discussion

LP-Micro leverages polynomial group lasso, ML algorithms and permutation importance tests to improve incident outcome (e.g., future disease) prediction using longitudinal microbial data. Compared to other ML/DL algorithms, LP-Micro addresses the challenges of leveraging the high-dimensional and longitudinal microbiome data for disease prediction or early detection. As shown by our simulation, popular ML/DL algorithms such as LSTM and its variants tend to overfit microbial data, resulting in poor prediction accuracy. Conversely, by introducing feature pre-screening and ensembling techniques, we demonstrate that ML/DL algorithms become more powerful in capturing the complicated time-varying association between microbial features and disease outcomes. Additionally, LP-Micro provides more interpretable association results in this context. Compared to the traditional generalized linear models, the ML algorithms in LP-Micro avoid making parametric assumptions regarding microbiome-disease associations so that it typically provides higher prediction accuracy and better flexibility to be suitable the data distribution and the non-linear association. Moreover, LP-Micro uses permutation-based tests that can better accommodate the skewness and zero-inflation of microbial data compared to HRT.

LP-Micro has shown favorable performance in both simulations and real data. In the simulations, LP-Micro outperforms the other state of the art ML methods in terms of identifying the right set of features and accuracy or prediction. In the VicGen study to predict ECC, LP-Micro is among the two best cumulative prediction methods and the only method that can provide *p*-values of important features for interpretation.

Strikingly, the significant important taxa output from LP-Micro identifies *S. mutans* abundance as highly predictive of future disease occurrence, confirming previous studies [12, 61]. LP-Micro extends this observation as the visit-wise analysis results also suggested the 5th visit at about 39 months of age is most important for prediction of ECC at age 5 (Figure 5a, 6a, 6b). In addition, and consistent with the polymicrobial aetiology of ECC, increased relative abundances of *S. vestibularis* and *S. wiggsiae*, that have both previously been associated with disease [64, 65], were identified as being significantly predictive of ECC at five years of age in our analysis. A decrease in relative abundance of *P. pasteri* was also identified as a novel biomarker of future disease, which is consistent with current understanding of ECC microbial aetiology and the pH sensitivity of this species. In the weight loss study, the pre-existing clinical/demographic variables, especially ages, are important for prediction of BMI change at the end of year one (Figure 7b). Specifically, older ages are associated with worse BS results, i.e., less BMI decrease (Supplementary Figure 8). Among the microbial biomarkers, *Schaalia* and *Cobetia* emerged as particularly significant, demonstrating their highest importance during the early and middle time points, while *Litorilituus* became more prominent at the one-year mark (Figures 5d, 7a, 7b). *Schaalia*, formerly *Actinomyces* [66], potentially due to its association with host infection, has previously been linked to weight loss [67]. This finding aligns with our observation that elevated levels of *Schaalia* correlate with weight loss (Supplementary Figure 8). *Cobetia* has been shown to be positively associated with the metabolite 5-hydroxytryptophan (5-HTP) [68], which has been used in the treatment of obesity [69]. Metabolite 5-HTP is a precursor to serotonin and has been studied for its role in regulating appetite and mood, both of which are critical in obesity management [70]. By increasing serotonin levels, 5-HTP may reduce appetite and cravings, particularly for carbohydrates, which in turn supports weight management efforts [71]. These prior findings provide further support for the results observed in our study concerning these two taxa. Notably, *Cobetia* exhibited significant relevance pre-surgery, suggesting its potential as a biomarker for predicting patient response to BS. Furthermore, our analysis revealed that, of the two surgery types examined, RYGB was associated with superior outcomes, a finding supported by previous research [72].

In this paper, we demonstrate the predictive performance of LP-Micro for four popular ML algorithms: NN, SVM, XGBoost and RF. However, the LP-Micro framework is highly flexible and can be extended to more advanced ML/DL algorithms. For example, in our simulation, LP-Micro improves the performance of GRU and CNN-GRU. Although such models are inferior to other ML algorithms in our simulation, they may gain in power in larger sample sizes. In conclusion, we provide comprehensive evidence from simulations and applications to two clinical datasets that the new method, LP-Micro offers advantages over existing approaches for powerful outcome prediction and feature interpretation in the context of time-varying microbiome data. Furthermore, while LP-Micro currently requires aligned time points of measurement, it can be extended to accommodate irregular longitudinal microbiome data in the future. For example, the prediction of using gut microbiome data of the Inflammatory Bowel Disease (IBD) patients at different time points [15] may be a different type of prediction since the patients don’t start from the same time points of disease progression. However, the prediction of the treatment effect may still be desirable. We used two well-designed microbiome studies that had samples from the same subjects at multiple time points to reflect the progression of the childhood oral microbiome and its association with ECC, or the longitudinal microbial changes after BS and how they are associated with the weight loss (measured as decreased BMI). There are currently few available longitudinal microbiome datasets that have both clear starting time points, e.g., two months of age or BS, and clear well-defined outcomes after a period, e.g., ECC or weight loss. However, much more of such data will become available in the future using sequencing techniques that can provide higher sequencing resolution than 16s rRNA gene sequencing. LP-Micro will be a valuable tool for the analyses of these cohorts.

## Methods

### Description of the two longitudinal microbiome datasets

The first dataset, derived from the VicGen longitudinal cohort established in 2008, aimed to study the natural history and causal factors of Early Childhood Caries (ECC) in infants and children [12, 52, 53]. Participants were recruited through their mothers from Maternal and Child Health Centers in six local government areas in Victoria, Australia. Clinical oral examinations and collection of saliva from the children were conducted at seven time points (mean ages ± standard deviation): 1.9 ± 0.8, 7.7 ± 1.3, 13.2 ± 1.2, 19.7 ± 2.0, 39.0 ± 3.2, 48.6 ± 1.6 and 60 ± 1.8 months [12, 52, 53]. Saliva samples collected at the first six visits were analyzed by 16s rRNA gene sequencing of the V4 region, and a total of 356 OTUs were identified. Of the 134 children whose oral microbiome was determined in this cohort, 69 remained healthy (i.e., cavity-free) throughout the study. By 39 months-of-age (the 5th visit), 12 children had a dental cavitation (defined as by having at least one ICDAS score of 3 or higher, [56]). By the time of the 6th visit (48.6 months-of-age), the number of children with at least one dental cavitation rose to 45. At the study’s conclusion, 65 children had developed cavities by 60 months-of-age. Our analysis aimed to predict children’s final ECC status using compositional microbial abundance data from the first six visits. We corrected anomalous samples that presented with visit times of the 5th visit later than that of the 6th visit. There are 100 subjects who have all the first six time points up to about 48 months, without any missing time points. We include data from these 100 subjects in our analysis, in which 42 subjects had ECC at about 60 months.

The second dataset records patients’ weight loss following BS. The cohort recruited 144 participants undergoing BS (50% Roux-en-Y Gastric Bypass (RYGB) and 50% Sleeve Gastrectomy (SG)) at the beginning of the study. These surgeries involve significant anatomical and physiological alterations that lead to changes in behavior and biology [54]. Body mass index (BMI) measurements and fecal material were collected from individuals at 1, 6, 12, 18 and 24 months post-surgery. Participants’ microbial profiles in the BS study were characterized through shotgun whole genome sequencing for time-series metagenomics data of 120 subjects (*n* = 120, [62]). A total of 1,533 microbial taxa were identified. Participants’ demographic data were collected before surgery, including age, race, height, and sex. Due to the high dropout rates (*>*50%) beyond 12 months, our analysis focused on predicting BMI changes at 12 months post-surgery, using log-normalized microbial genera counts, where 35 patients with incomplete visits were excluded. There are 84 subjects who have all four time points at pre-surgery, 1, 6, 12, without any missing time points up to 12 months. We include data from these 84 subjects in our analysis.

### Polynomial group lasso for longitudinal feature selection

Let **x**_*i*_ = (*x*_*i*1_, …, *x*_*i*(*p×q*)_)^*T*^ be a *p×q*-dimensional concatenated vector of the abundance of *p* microbial taxa measured at *q* time points from the *i*-th subject. Let *y*_*i*_ be the outcome variable to be predicted. For a cohort with *n* samples, define the design matrix **X** = [**x**_1_, …, **x**_*n*_]^*T*^ and the response vector **y** = (*y*_1_, …, *y*_*n*_)^*T*^. The objective is to identify a subset of microbial taxa that are temporally associated with the clinical outcomes.

To facilitate this, we employ the group lasso estimator, an extension of the lasso regularization technique [48, 49]. The group lasso estimator is defined as:

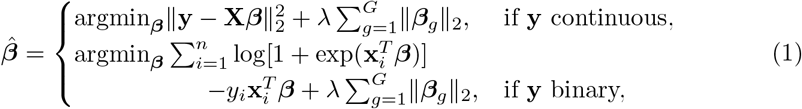

where ***β*** represents the vector of coefficients, *G* is the total number of predefined variable groups, *λ* is the regularization parameter, ***β***_*g*_ denotes the indices corresponding to the *g*-th group, and ∥*·*∥_2_ denotes the ℓ_2_ norm. Unlike the standard lasso estimator, group lasso promotes sparsity across predefined groups of coefficients rather than individual coefficients. This characteristic is beneficial for screening high-dimensional variables that can be naturally clustered into coherent groups. In longitudinal data analysis, these groups can be defined by the repeated measurements of a microbial taxon. Thus, group lasso preserves the entire temporal trajectory of these measurements if they show significant correlation with clinical outcomes. In contrast, the standard lasso might only select sporadic points from this trajectory, potentially overlooking important patterns.

One major concern of utilizing group lasso on longitudinal microbiome data is its assumption of linear effects. Due to the linear constraint on the functional forms, the variable selection process may suffer from being unstable. For example, a simple transformation of the raw count may alter the selection of microbiome taxa. Inspired by previous work [45, 46], we employ natural splines based on the scaled data to approximate more general functional forms of the original data to relax the linear problem assumptions in group lasso. Specifically, we define **z**_*ij*_ = (*ϕ*_1_(*x*_*ij*_), …, *ϕ*_*M*_ (*x*_*ij*_))^*T*^, where *ϕ*_*m*_(*·*) represents the spline function of order *m*. Thus, the polynomial design matrix of the *j*-th covariate is **Z**_*j*_ = [**z**_1*j*_, …, **z**_*nj*_]^*T*^, and the total polynomial design matrix is **Z** = [**Z**_1_, …, **Z**_*j*_]. In the polynomial group lasso for longitudinal data, we replace **X** by **Z** in the following modified group lasso equation (1):

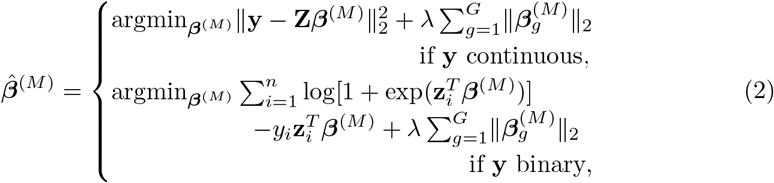

where *M* is a predefined order to approximate the non-linear functions. In this paper, we set *M* to 3.

To be noted, if there are other pre-existing clinical or demographic variables reported before (or at) the first microbiome-collection time point, that are needed to be considered in prediction, **x**_*i*_ in design matrix will be accordingly modified to include these variables to be screened together with the other longitudinal features in polynomial group lasso. In addition, it’s also possible that there are other longitudinal clinical or demographic variables to be used for prediction, that will be considered together with the longitudinal microbial features by modification of the design matrix.

### Visit-wise and cumulative longitudinal prediction

We train various ML models to predict the clinical outcomes from the chosen taxa from microbiome data. Details on the ML algorithms employed will be provided in a subsequent section. The prediction strategies include: (a) visit-wise prediction, utilizing microbiome profiles from one visit, and (b) cumulative prediction, incorporating all microbiome profiles up to a chosen visit. Specifically, the visit-wise prediction models at the *k*-th visit utilize microbiome data from only that time point to predict clinical outcomes, i.e., estimating 𝔼 [*y*|**X**_*·*(*p×k*+1:*p×*(*k*+1))_], where **X**_*·*(*a*:*b*)_ denotes columns indexed from *a* to *b*− 1. In contrast, the cumulative prediction models at the *k*-th visit estimate 𝔼 [*y*|**X**_*·*(1:*p×*(*k*+1))_]. The fitted models at the *k*-th visit are denoted as 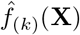 for visit-wise, and 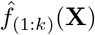 for cumulative predictions. The features used for cumulative prediction at this step are the selected microbial taxa by the above procedure of polynomial group lasso.

To determine at which visit the microbial abundance provides the most accurate prediction of outcomes, we evaluate the *q* cumulative models on an independent testing set. The difference in performance metrics between the models for the (*k* − 1)-th and *k*-th visits indicates the additional information in prediction from a visit. For instance, using MSE to assess the model performance, the increment in model prediction can be quantified by

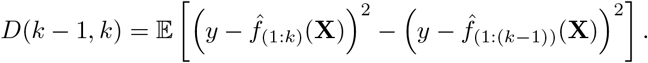

A negative or zero value of *D*(*k* − 1, *k*) suggests that the *k*-th repeated measurement of microbiome levels does not enhance disease prediction. In practice, the information increment *D*(*k* − 1, *k*) is estimated by 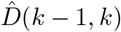,or

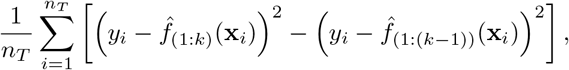

where *n*_*T*_ denotes the sample size of the testing set. Upon identifying the optimal visit for prediction by each ML algorithm, we rank these algorithms based on their performance metrics at their respective optimal visits.

### Feature importance score for explainable longitudinal predictors

To understand the time-varying association between microbiome profiles and clinical outcomes, we need to investigate three types of feature effects: (i) variable-wise effect, which examines the impact of a microbial taxon on the disease outcome at each single time point, (ii) taxon-wise effect, which calculates the average impact between a microbial taxon and the disease outcome across multiple time points, (iii) visit-wise effect, which assesses the average impact of all selected microbial traits at a single time point. LP-Micro provides an importance score and a corresponding p-value for each of these effects. These metrics are derived from top-ranked ML models 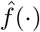 from the previous section. In this paper, we utilize the best two models to evaluate the microbial effects.

Inspired by previous studies [38, 42], we quantify the three types of effects defined above by importance scores. For each type of effect, we replace the corresponding columns of the previously defined design matrix **X**, where columns represent the *p × q* features by combination of *p* taxa and *q* visits, by generating replicates from the same distribution but breaking the association with the disease outcome, denoted as **X**^*′′*^. Specifically, for each variable-wise effect, we replace one column with its permutation replicates, producing *p × q* importance scores and *p*-values, while for each taxa-wise or visit-wise effect, *q* or *p* columns corresponding to the information from the taxon or visit are replaced, producing *p* or *q* importance scores and *p*-values, respectively. Similarly, as in feature selection, **x**_*i*_ in design matrix **X** can be modified here for importance score if other pre-existing clinical or demographic variables, as well as other clinical predictors, need to be considered in prediction.

We calculate those importance scores and *p*-values by evaluating the performance of the ML model 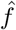 with the permuted design matrix **X**^*′′*^ as input using the following empirical loss functions:

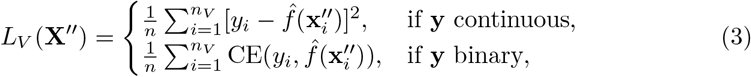

where CE(*y, ŷ*) = −*y* log *ŷ*−(1−*y*) log(1−*ŷ*), the cross-entropy loss for binary response, and *n*_*V*_ is the sample size of an independent validation set *V*. Under the null hypothesis that an effect is not predictive, the expected mean difference in loss, 𝔼 [*M*_*V*_ (**X**^*′′*^)] = 𝔼 [*L*_*V*_ (**X**^*′′*^) − *L*_*V*_ (**X**)] = 0, should equal zero. This mean difference is termed the effect importance score. To accommodate the skewed microbial distribution with excessive zeroes, we employ group permutation instead of multivariate normal bootstrapping to generate null samples, as inspired by previous works [40, 42].

To accurately estimate the feature importance score, we adopt the *K*-fold cross-fitting strategy. Suppose we split the data into *C* distinct validation sets, denoted as *V*_1_, …, *V*_*C*_. For each validation set, we fit an ML model using the rest of the data. The cross-validated effect importance score *M*_(*CV*)_(**X**^*′′*^) can be computed by taking the arithmetic average of 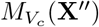 with respect to their sample size, where *c* = 1, …, *C. M*_(*CV*)_(**X**^*′′*^) is defined as the feature importance score. Its one-sided *p*-value is then computed by assuming normality of *M*_(*CV*)_(**X**^*′′*^), and the empirical variance 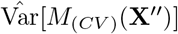 can be computed by taking the arithmetic average of:

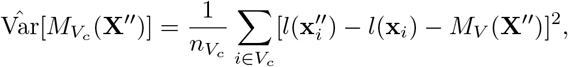

where *l*(**x**_*i*_) (or 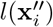) denotes the value of mean squared loss or cross-entropy loss defined in equation (3) evaluated at the *i*-th sample in the validation set (or its permuted version), and 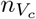 denotes the sample size of the validation set *V*_*c*_.

### Machine learning algorithms in LP-Micro

SVM is implemented via R package “e1071” using Radial kernels. RF and XGBoost are implemented via R package “randomForest” and “xgboost” respectively. Their hyperparameters are identified via five-fold cross-validation.

### Deep learning architectures in LP-Micro

LSTM, GRU, and CNN-GRU are implemented via Keras and TensorFlow for R. Fully connected deep neural network is implemented via R package “deepTL”. The detailed parameter settings are provided in a Supplementary section. To stabilize the performance of deep architectures, LP-Micro includes the following ensemble mechanism [43]: (i) we first train *B* deep learning models via bootstrapping training samples, denoted as 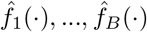,(ii) next we evaluate trained models on out-of-bag (OOB) sample sets *O*_1_, …, *O*_*B*_. The final models are obtained by aggregating the models: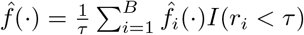, where *r*_*i*_ is the rank of model performance. We set *B* to 50 and *τ* to 40 for all deep learning algorithms.

### Simulation design

We evaluate the performance of LP-Micro with the following data generation model. Specifically, we simulate the continuous disease outcome *y*_*i*_ as

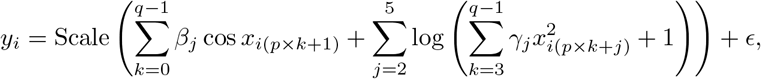

where **x**_*i*_ = (*x*_*i*1_, …, *x*_*i*(*p×q*)_)^*T*^ represents *p* microbial taxa abundance collected from *q* repeated visits, following Poisson log-normal distribution *PN* (*µ*, **Σ**), and *β*_*j*_ and *γ*_*j*_ are coefficients sampled from uniform distributions 𝒰 (−2, 2) and 𝒰 (0, 4) at the beginning of each simulation and fixed afterwards. This model assumes one microbial taxon is effective across all five visits, while the other four taxa are only effective in the last two visits, with 13 nonlinear covariates in total. To mimic the highly correlated microbial data, we pose within-visit correlation to **x**_*i*_ by setting its covariance **Σ** = diag(**Σ**_1_, …, **Σ**_*q*_), where **Σ**_*j*_ is a *p×p* matrix with diagonal entries of 1 and off-diagonal entries of 0.1. After normalizing the response for all covariates to zero mean and unit variance, noise *ϵ* following the normal distribution 𝒩 (0, 0.1) is added.

We set both the training and testing sample size to *n* = 120 and the number of repeated measurement *q* set to 5. To investigate the impact of the number of microbial taxa, we vary the number of microbial taxa *p* across *{*100, 200, 500*}*, simulating three levels of sparsity (5%, 2.5%, 1%). Each scenario is randomly repeated for 50 times, where the hyper-parameter of group lasso is set at half of the largest *λ* that allows non-zero coefficients, and 5-fold cross-validation is used to compute feature importance scores and *p*-values.

### Real data analysis

The ECC and weight loss data are split into a training set, a validation set, and a testing set. Specifically, 75% of the samples are allocated to the training set for variable selection and model training. The validation set comprises 10 samples from the remaining 25% to select the group lasso hyperparameter. The remaining samples constitute the testing set used to evaluate the models.

To evaluate prediction results, for the ECC data, we compute accuracy and AUC, and for the weight loss data, we compute MSE and PCC. In accordance with previous studies [12], we categorize ECC data into two groups for group lasso: the first four visits and the last two visits. For baseline models, we fit either a logistic regression model or a linear regression model with an *l*_1_ penalty. The penalty parameter is selected via 10-fold cross-validation. This entire procedure is repeated 20 times.

## Supplementary information

The supplementary document can be found online. This study was supported by national institutes of health 1R03DE034507-01 and U01DE025046 grant. The authors would like to acknowledge all the families who gave up time to be involved in the VicGen cohort, the Maternal and Child Health Nurses and the administration staff who assisted with recruitment.

## Code availability

The implementation of LP-Micro can be found in a GitHub repository (https://github.com/IV012/LPMicro).

## Data availability

The metagenomic sequences of the weight loss study can be found at the National Center for Biotechnology information Sequence Read Archive (https://www.ncbi.nlm.nih.gov/sra) under BiopProject PRJNA668357 and PRJNA668472. VicGen cohort data are available upon request by contacting the team through its project website (https://mspgh.unimelb.edu.au/centres-institutes/centre-for-health-equity/research-group/child-community-wellbeing/research/previous-projects/vicgen).

