## Supplementary file for "Longitudinal Microbiome-based Interpretable Machine Learning for Identification of Time-Varying Biomarkers in Early Prediction of Disease Outcomes"

### Exploring Different Deep Learning Architectures in Simulation Studies

To explore the impact of the architectures of deep learning (DL) models, we implemented fully-connected deep neural networks (NN), long short-term memory (LSTM [1, 2, 3]), gated recurrent units (GRU [4, 5]), and convolutional neural network (CNN)-GRU. The hidden states of LSTM and GRU were set to (256, 256). For CNN-GRU, the number of filters and the kernel size the of CNN were set to 32 and 2, respectively, followed by two layers of GRU with hidden states set to (256, 128). Between each layer, a dropout layer with dropout rate 0.5 was included to avoid overfitting. The activation function was chosen as rectified linear unit. We used Adam [6] as the optimizer with the learning rate set to 0.001. The prediction results of those models are summarized in Supplementary Table 1, which exhibit minor improvement of using more advanced strutures than simple NN. LP-Micro, in contrast, significantly improved the prediction accuracy of all deep learning models. Specifically, LP-Micro NN and CNN-GRU outperformed other DL models.

| Pipelines | p=100 | | p=200 | | p=500 | |
|---|---|---|---|---|---|---|
| | **MSE** | **PCC** | **MSE** | **PCC** | **MSE** | **PCC** |
| **Benchmarks** | | | | | | |
| NN | 1.05 | 0.10 | 1.02 | 0.10 | 1.00 | 0.10 |
| LSTM | 1.10 | 0.02 | 1.09 | 0.02 | 1.09 | 0.03 |
| GRU | 1.25 | 0.15 | 1.22 | 0.12 | 1.11 | 0.10 |
| CNN-GRU | 1.04 | 0.22 | 1.02 | 0.21 | 1.01 | 0.15 |
| **LP-Micro** | | | | | | |
| NN | 0.61 | **0.64** | **0.64** | 0.61 | 0.73 | 0.52 |
| LSTM | 0.96 | 0.37 | 1.01 | 0.29 | 1.00 | 0.17 |
| GRU | 0.72 | 0.57 | 0.71 | 0.57 | 0.89 | 0.36 |
| CNN-GRU | **0.60** | 0.62 | 0.65 | **0.62** | **0.72** | **0.53** |

Supplementary Table 1: **Performance of deep architectures in simulation.**

### Comparison between Hold-out Randomized Tests and Permutation Tests

In order to demonstrate the extra power using permutation test compared to other parametric tests, we replace the permutation importance score in LP-Micro with hold-out randomized tests (HRTs, [7]), which assume that the predictors (i.e., microbial abundance) are normally distributed. As shown by Supplementary Figure 1, the performance of HRTs is inferior to the corresponding permuting methods in Figure 2a if the data distribution is non-Gaussian. Specifically, although LP-Micro can still identify the temporal-microbial patterns from the simulated data (i.e., one taxon is important at five time points, while the other four taxa are important at the last two time points), the HRT importance scores of these important features are less distant from the unimportant features than those of permutation importance scores that we utilized for real data analysis.

### Analysis of VicGen Cohort for Early Childhood Caries (ECC)

Supplementary Figure 2 demonstrates the sample similarities using the microbial abundance data either of all taxa or of the significant taxa based on feature importance in the interpretation step of LP-Micro. This tSNE figure suggests that the microbial abundance of children in VicGen cohort is highly time-varying, of which the characteristic is preserved by features identified by LP-Micro.

Supplementary Figure 3 and 4 present the longitudinal distribution of microbial taxa identified by LP-Micro. Considering the zero-inflated nature of microbial distributions, we performed differential prevalence

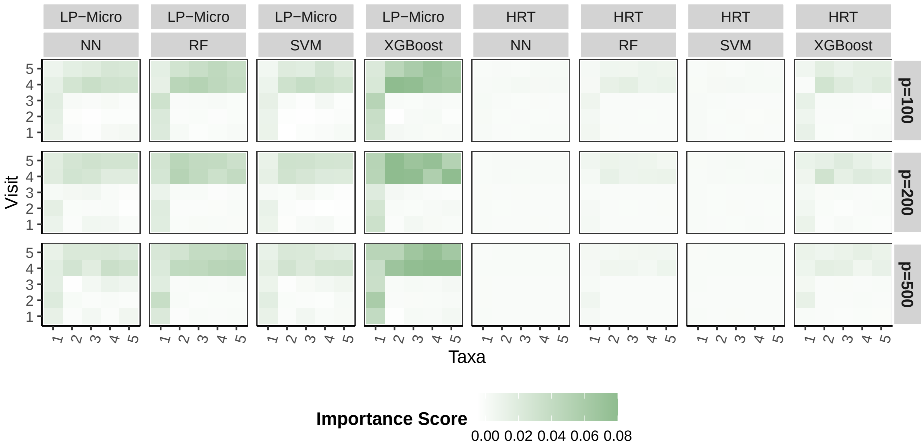


Supplementary Figure 1: **Feature importance in simulations using Hold-Out Randomized Tests (HRT) and LP-Micro based on HRT.** Microbial feature importance at each time point. The x-axis represents five causal microbial taxa, and the y-axis represents each of five visits. Each grid represents the feature importance of the microbial taxa during the corresponding visit row.

and abundance analysis of those identified taxa between children with and without cavities. Four microbial taxa, including *Lachnospiraceae [G-2]* oral taxon 096, *Megasphaera micronuciformis*, *Pedobacter* oral taxon 321, and *Streptococcus mutans*, are more prevalent among children with cavities across $\geq 5$ time points out of all six time points. Additionally, despite the low prevalence of *Scardovia wiggsiae* in the cohort in most time points, it is highly predictive of ECC at the last two visits, exhibiting a difference of $> 10\%$ in prevalence between health and disease groups. Moreover, five identified microbial taxa with high prevalence in both groups exhibit the same direction in their abundance effects across $\geq 5$ out of all 6 time points, including *S. mutans*, *Porphyromonas pasteri*, *Streptococcus vestibularis*, *Lachnospiraceae [G-2]* oral taxon 096, and *Haemophilus parainfluenzae.* Interestingly, higher abundance of *Lachnospiraceae [G-2]* oral taxon 096 and *Porphyromonas pasteri* seems to be inversely associated with ECC.

### Analysis of the Weight Loss Study following Bariatric Surgery (BS)

Because the ability of baseline microbial profiles to predict weight loss after BS was limited in this study, we also implemented LP-Micro without the baseline microbiome information (i.e., retaining microbiome information only after surgery, see Supplementary Figures 5, 6, and 7). Here, LP-Micro still has superior performance than the standard ML models using all taxa (as the baseline model) and the sPLS-based prediction using sPLS selected features (Supplementary Figure 5), same as in Figure 4b. LP-Micro, as a

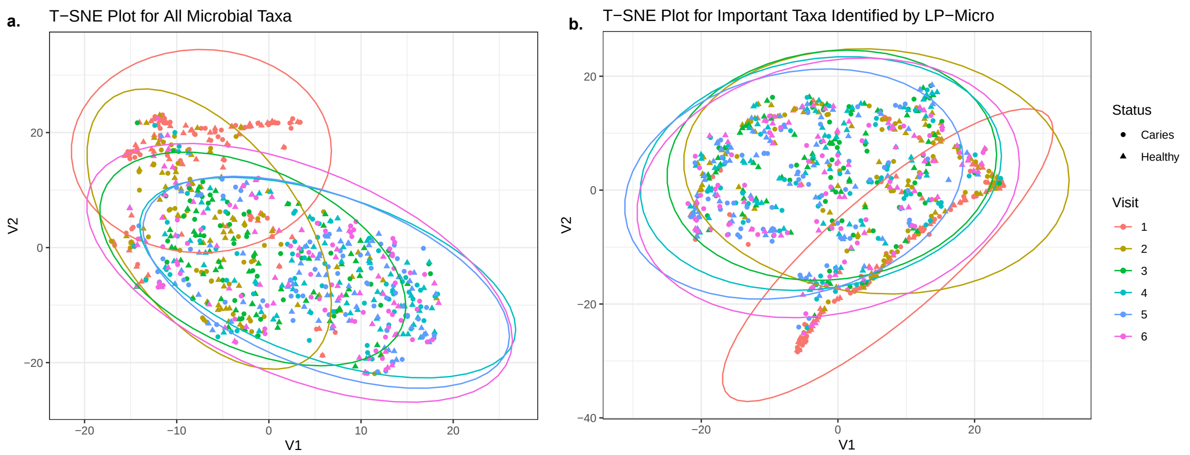


Supplementary Figure 2: **Temporal dynamics of oral microbial level for children in VicGen cohort visualized by t-distributed Stochastic Neighbor Embedding (tSNE). (a)** Dynamics of all microbial taxa in the study. **(b)** Dynamics of important microbial taxa identified by LP-Micro. Each dot is for a sample. Features used in (b) either have p-value of PermFit-RF LP-Micro or PermFIT-XGB in LP-Micro $< 0.1$, i.e., the significant taxa in Figure 6b.

powerful approach for cumulative prediction, also strongly preserves superior performance when comparing to the visit-wise and other cumulative prediction (Supplementary Figure 6a), same as in Figure 5c. Specifically, without LP-Micro, incorporating longitudinal microbial information may not be beneficial for prediction. For instance, the prediction PCC of SVM is higher than 0.4 with only low-dimensional demographic variables collected prior to surgery, and the PCC drops to <0.2 when incorporating the high-dimensional microbial features collected at 1 or 12 months post-surgery. Its performance becomes even worse when using cumulative microbial information. For instance, SVM outperforms other models with microbial information in visit-wise prediction at 6 months post-surgery. However, its prediction becomes worse than lasso and NN when adding information at one month post-surgery, with a significant decrease in prediction PCC. In contrast, the performances of SVM are more stable within our proposed framework, as indicated by the LP-Micro plot in Supplementary Figure 6a.

Supplementary Figure 8 shows LP-Micro's capability to discriminate BMI change using longitudinal microbiome data. The observed more variation across multiple time points in Supplementary Figure 8b than in Supplementary Figure 8a suggests that high capability to discriminate the BMI change across time points in the significant taxa obtained from the third step of LP-Micro for interpretation, therefore LP-Micro can select predictive microbial biomarkers for weight loss. Supplementary Figure 9 further presents

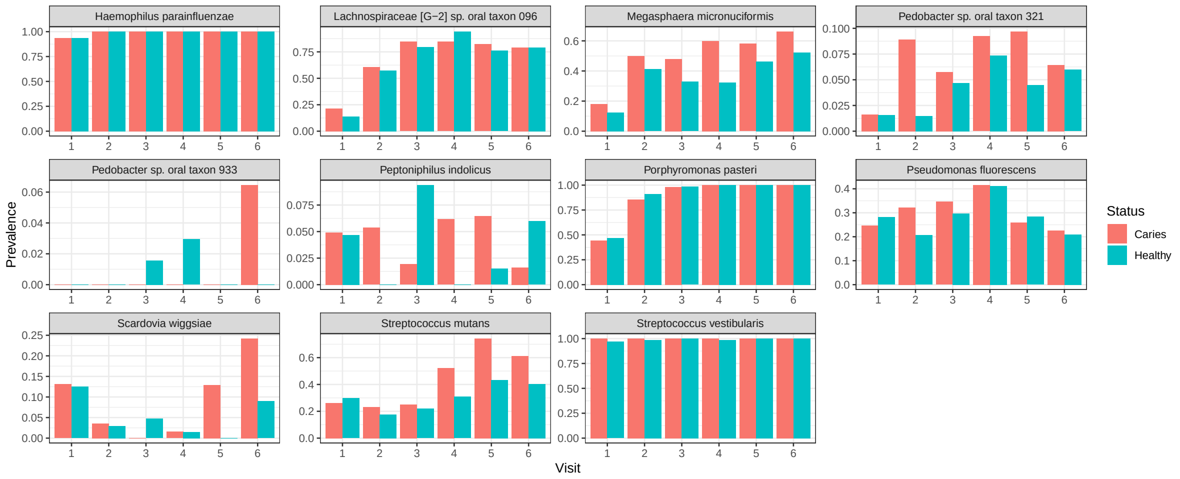


Supplementary Figure 3: **Prevalence of important taxa identified by LP-Micro in the VicGen ECC cohort for children.** The x-axis corresponds to the six study visits, while the y-axis indicates the prevalence, defined as the proportion of samples where a taxon shows detectable abundance (i.e., nonzero abundance) at each time point. The presented taxa correspond to those identified as significant in Figure 6b.

the association between LP-Micro selected features (based on $p$-values in the third step) and BMI change for the 12-month period between pre-surgery and post-surgery. Despite the relatively linear trend between age and BMI reduction, other biomarkers show non-linear associations with BMI change (Supplementary Figure 9).

[4] Kyunghyun Cho, Bart van Merrienboer, Caglar Gulcehre, Dzmitry Bahdanau, Fethi Bougares, Holger Schwenk, and Yoshua Bengio. Learning phrase representations using rnn encoder-decoder for statistical

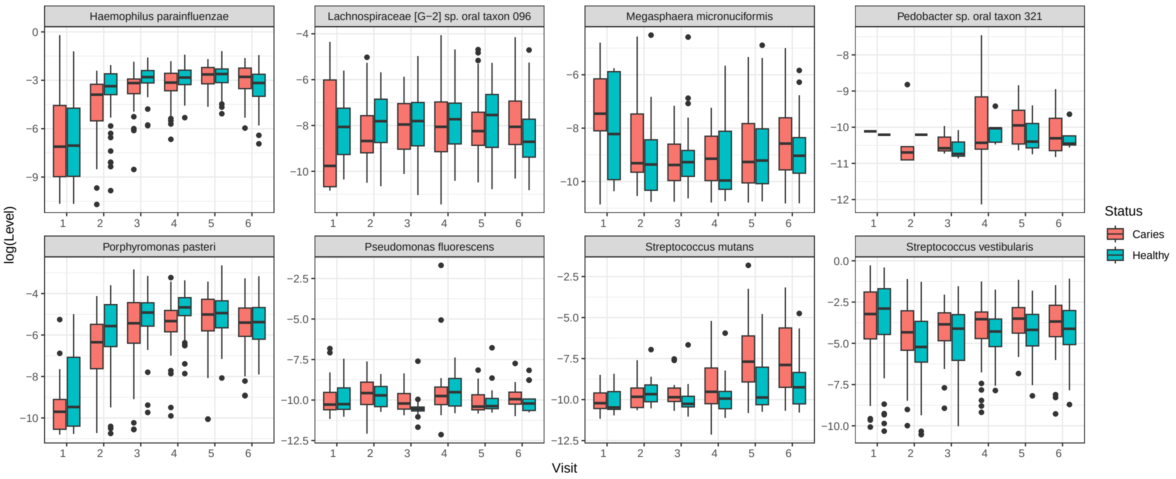


Supplementary Figure 4: **The distribution of log-abundance of important taxa identified by LP-Micro for children in VicGen ECC cohort.** The x-axis represents the six visits of the study, and the y-axis represents the log-abundance level at each visit time point in which natural logarithm was used. Samples with zero abundance of the corresponding taxa are not included in the figure. Here the eight taxa are selected among the 11 significant taxa in Figure 6b if they have relative less number of zeros and higher non-zero abundance.

machine translation. In Proceedings of the 2014 Conference on Empirical Methods in Natural Language Processing (EMNLP), pages 1724–1734. Association for Computational Linguistics, 2014.

[5] Xingjian Chen, Lingjing Liu, Weitong Zhang, Jianyi Yang, and Ka-Chun Wong. Human host status inference from temporal microbiome changes via recurrent neural networks. Briefings in Bioinformatics, 22(6):bbab223, 06 2021.

[6] Diederik P Kingma. Adam: A method for stochastic optimization. arXiv preprint arXiv:1412.6980, 2014.

[7] Wesley Tansey, Victor Veitch, Haoran Zhang, Raul Rabadan, and David M. Blei. The holdout randomization test for feature selection in black box models. Journal of Computational and Graphical Statistics, 31(1):151–162, 2022.

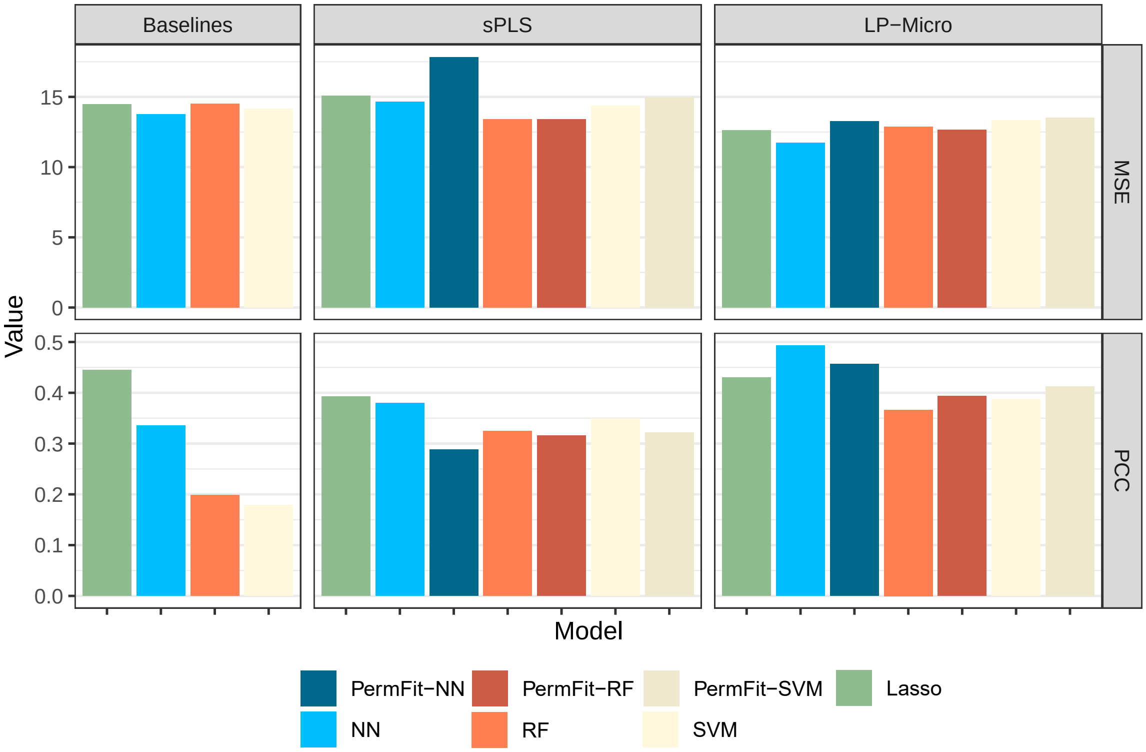


Supplementary Figure 5: **Prediction results of machine learning models in the weight loss study using baseline features, sPLS features, and LP-Micro (group lasso) features on patients' demographics and microbial profiles one month post-surgery.** The average mean squared error (MSE) and Pearson correlation (PCC) between the body mass index (BMI) change in 12 months and the predicted outcome.

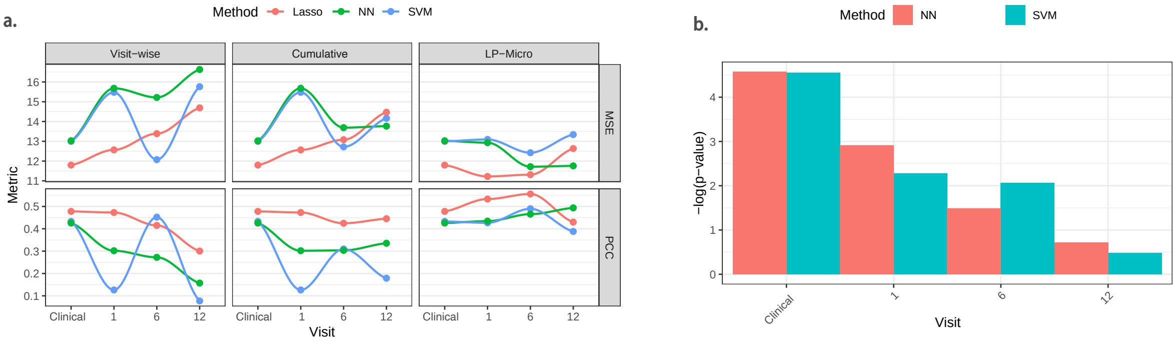


Supplementary Figure 6: **Results of visit-wise prediction, cumulative prediction, and LP-Micro to investigate weight loss after bariatric surgery.** To accomplish visit-wise prediction and cumulative prediction, ML algorithms are trained using all microbial features. LP-Micro refines the cumulative prediction by microbial feature pre-screening and model ensembling. The pre-surgery microbial profiles are removed in all methods, while demographic variables are kept, labeled as 'clinical'. **(a) Prediction MSE and PCC of BMI change at each visit in the weight loss cohort. (b) Visit-wise microbial $p$-values of importance scores for BMI change prediction.** Visit importance scores are calculated using the LP-Micro models with microbial features up to 12 months after BS in (b).

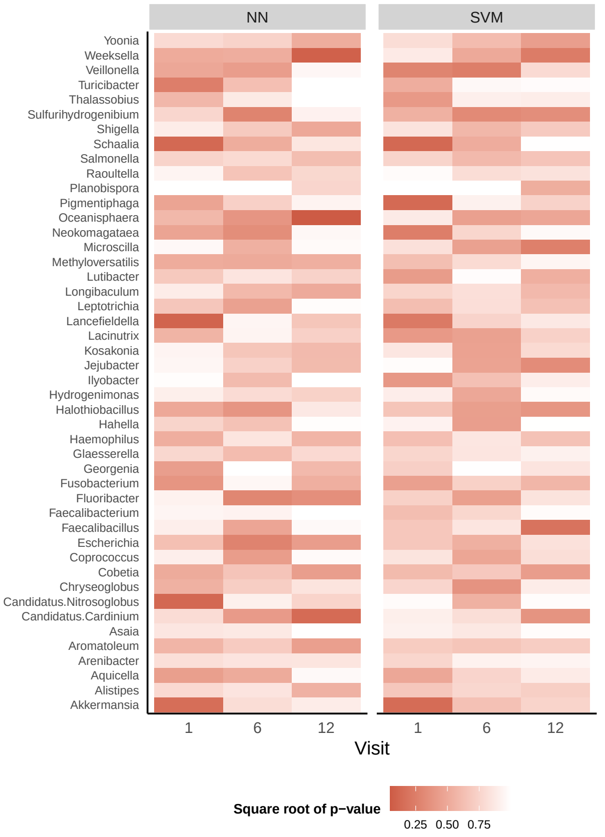


(a) Point-wise feature importance in weight loss data.

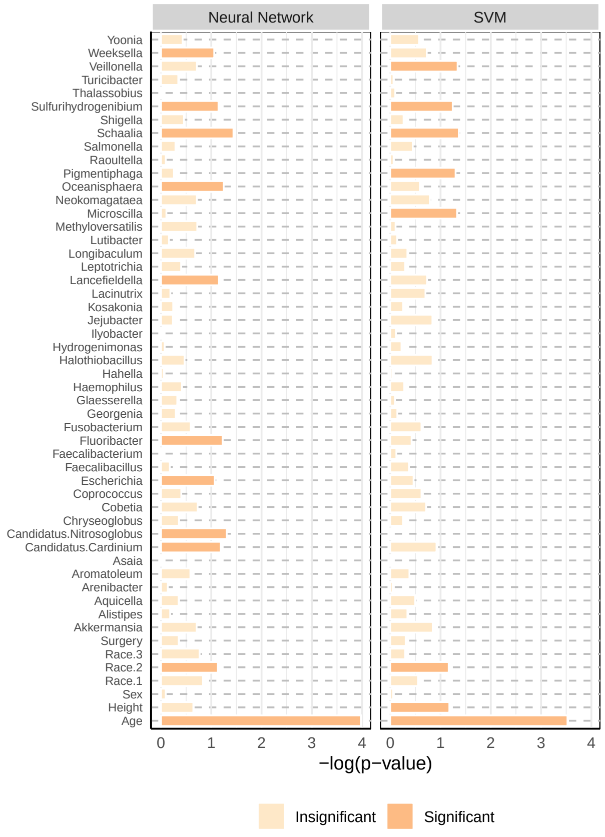


(b) Taxon-wise negative $\log_{10}$ p-values in weight loss data.

Supplementary Figure 7: **Feature importance for the weight loss data (pre-surgery microbial profiles removed).** The y-axis consists of microbiome genera selected by LP-Micro, and $p$-values are computed by PermFit. **(a)** Each grid represents the significance of the microbiome at each corresponding visit. Darker colors indicates higher importance. **(b)** The x-axis represents the $\log_{10}$ p-values for each taxon in terms of their longitudinal effects on BMI change.

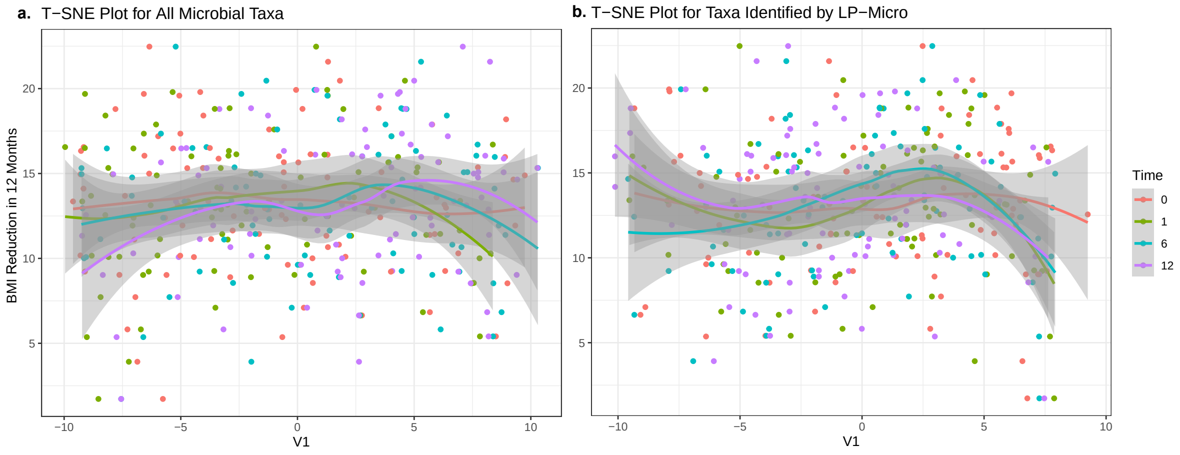


Supplementary Figure 8: **Temporal dynamics of gut microbial level for patients in weight loss cohort visualized by tSNE.** The x-axis represents the tSNE score and the y-axis represents the BMI change in 12 months. **(a)** Dynamics of all microbial taxa in the study. **(b)** Dynamics of important microbial taxa identified by LP-Micro.

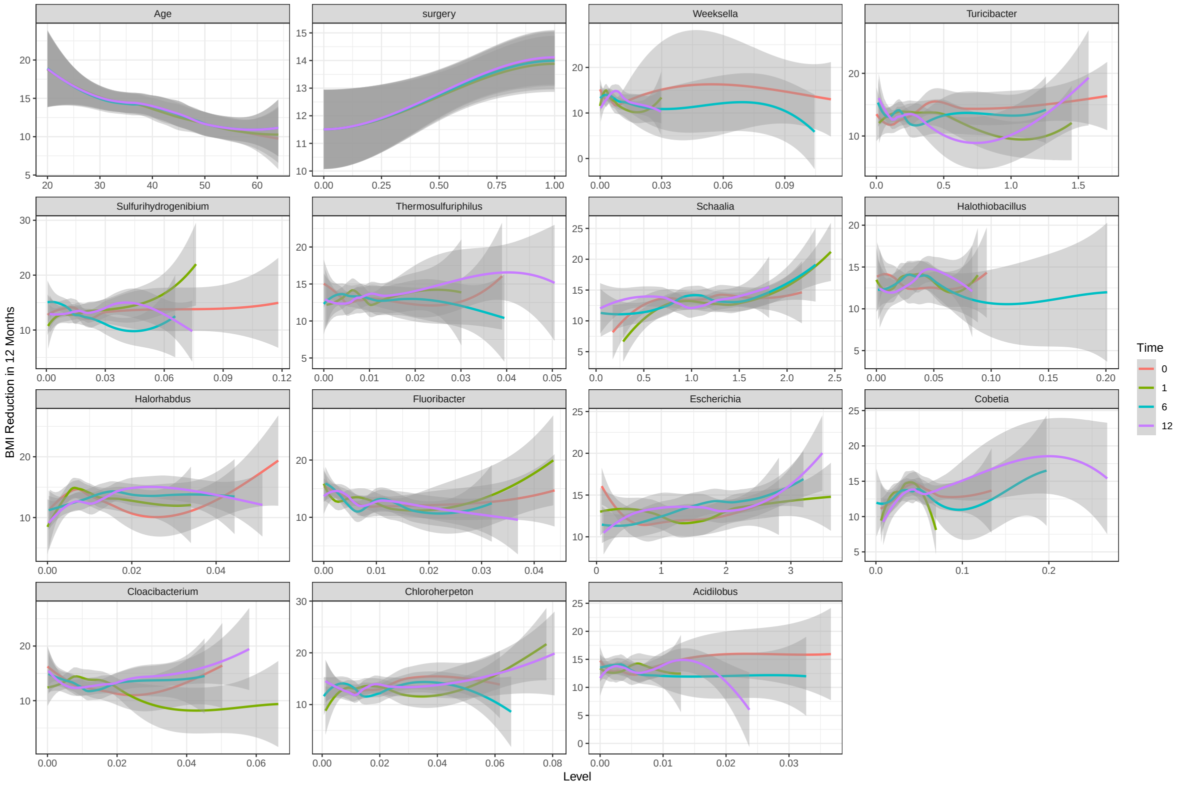


Supplementary Figure 9: **The association between BMI change after 12 months and levels of important features identified by LP-Micro for the weight loss study.** The x-axis represents the feature levels, and the y-axis represents the log-abundance level at the visit time.
